# Beta bursts in SMA mediate anticipatory muscle inhibition

**DOI:** 10.1101/2025.01.24.634650

**Authors:** Viktoriya Manyukhina, Oussama Abdoun, Franck Di Rienzo, Fanny Barlaam, Sébastien Daligault, Claude Delpuech, Maciej Szul, James Bonaiuto, Mathilde Bonnefond, Christina Schmitz

## Abstract

In motor networks, inhibition has been associated with oscillatory activity in the mu (8-12 Hz) or beta (13-30 Hz) bands, yet how these rhythms ultimately influence muscle activity remains unclear. The Bimanual Load-Lifting Task (BLLT) elicits anticipatory inhibition of the elbow flexors in the load-supporting arm during voluntary load-lifting, providing a suitable model to study oscillatory mechanisms of muscle inhibition. We recorded magnetoencephalography (MEG) in adult participants performing the BLLT. Optimal postural stabilization occurred when *Biceps brachii* inhibition preceded unloading by ∼10-40 ms. Stronger muscle inhibition within this time window was associated with reduced high-gamma (90-130 Hz) power, potentially reflecting reduced excitability, and increased high-beta power in the contralateral Supplementary Motor Area (SMA), with high-gamma power reduction partially mediating the effect of high-beta power on anticipatory inhibition. Furthermore, trials containing beta bursts (22-28 Hz) exhibited both optimally timed anticipatory inhibition and concurrent reduction in high-gamma power. These findings suggest that optimal anticipatory inhibition in the load-supporting elbow flexor is linked to reduced SMA excitability mediated by inhibitory beta bursts. More broadly, they provide new evidence linking beta bursts to the inhibitory control of muscle function, and contribute to a better understanding of motor anticipation.

## Introduction

Anticipatory postural adjustments (APA) are essential for smooth and coordinated motor behavior, as voluntary movements often cause postural perturbations that must be anticipated and compensated for (Massion, 1992). Yet, the neural mechanisms underlying APA, particularly those leading to the transient inhibition of tonic muscle activity, remain poorly understood. The bimanual load-lifting task (BLLT) provides a well-established paradigm to studying APA and their associated inhibitory muscle control in humans (Hugon et al., 1982). In this task, the postural arm supports a load placed on its wrist while the participant lifts it with the contralateral hand. Unloading induces a change in force, causing destabilization of the postural forearm, resulting in an upward movement. APA are initiated prior to the lifting of the object to counteract this perturbation, thereby stabilizing posture. A core signature of APA in the BLLT is the inhibition of postural elbow flexors, reflected in reduced electromyographic (EMG) activity. This anticipatory muscle inhibition precedes unloading and contributes to the horizontal forearm stabilization during load-lifting.

While prior research (Ng et al., 2013a) examined the network underlying APA in the BLLT, the specific mechanisms governing anticipatory muscle inhibition during this bimanual coordination – distinct from subsequent postural stabilization – remain unexplored. Previous studies have suggested that this muscle inhibition reflects an underlying cortical process, specifically the inhibition of contralateral cortical activity that ensures tonic muscle control for load maintenance, which is released in anticipation of load lifting (Kazennikov et al., 2005, 2006); however, this has not been directly demonstrated. A clear EMG signature of muscle inhibition also makes the BLLT a valuable framework for exploring the oscillatory mechanisms of motor inhibition, enabling the investigation of inhibitory control at the muscle level.

Previous studies investigating the oscillatory mechanisms of motor inhibition have reported mixed findings, with both mu (8-12 Hz) and beta (13-30 Hz) activity implicated in the inhibitory process. Elevated mu power has been associated with motor inhibition, particularly during action withholding and action observation tasks (Bönstrup et al., 2015; Köster and Meyer, 2023). Beta-band activity, predominantly expressed in bursts (Lundqvist et al., 2024), has been linked to movement slowing during the pre-movement phase (Khanna and Carmena, 2017; Little et al., 2019), and in the inhibition of initiated actions (Picazio et al., 2014; Jana et al., 2020; Schaum et al., 2020). However, the oscillatory mechanisms underlying anticipatory muscle inhibition remain unstudied, leaving it unclear whether mu- or beta-band activity mediates this process. Furthermore, it remains unclear whether these activities can contribute to inhibition at the level of the final motor output – the muscles – and what mechanisms underlie this effect. Here, we addressed these questions by examining the mechanisms of anticipatory muscle inhibition in the BLLT in adults. Our first aim was therefore to examine this relationship by linking mu and beta activity to muscle inhibition strength and cortical excitability, contributing to a fundamental understanding of how inhibitory control is implemented in the brain.

Motor inhibition has been consistently associated with decreased activity in the primary motor cortex (M1; Kazennikov et al., 2005, 2006; Borgomaneri et al., 2020). However, in the BLLT, Ng et al. (2013b) observed stronger beta desynchronization in the supplementary motor area (SMA) contralateral to the load-supporting forearm than in M1 prior to unloading, suggesting stronger activation of this area during APA. They therefore proposed that corticospinal projections originating from the SMA, rather than M1, may play a central role in APA regulation. To clarify this issue, we investigated whether anticipatory inhibition in the elbow flexors of the load-supporting forearm is associated with reduced activity in the contralateral M1 or SMA and examined the network supporting this inhibition. Specifically, we assessed the contribution of cortical and subcortical (basal ganglia and cerebellum) regions previously associated with voluntary unloading (Viallet et al., 1987; Schmitz et al., 2005; Ng et al., 2011, 2013b, 2013a). While the basal ganglia are critical for APA, the cerebellum has been proposed to ensure temporal coordination between postural and motor commands; however, their specific role in anticipatory inhibition of the postural elbow flexors remains unclear. Importantly, the BLLT is particularly well suited to address this question, as it allows postural stabilization processes to be dissociated from load lifting, as APA-related postural control and load-lifting motor control are largely segregated across hemispheres, consistent with findings that lesions contralateral, but not ipsilateral, to the postural forearm reduce APA (Massion, 1992; Viallet et al., 1992). Our second aim was therefore to explore the neural pathways underlying mature anticipatory muscle inhibition in adults, thereby advancing knowledge of the neural basis of motor anticipation.

## Methods

### Participants

Sixteen adults without neurological or psychiatric conditions (11 males, mean age 27 ± standard deviation (SD) 3.8 years) participated in this study. Handedness of the participants was determined using the Edinburgh Handedness Inventory (Oldfield, 1971). All participants were classified as right-handed. Written informed consent was obtained from each participant in accordance with the Declaration of Helsinki (2013 revision, Part 6). The study was approved by the French ethics committee (CPP Sud-Est III) and authorised by the French National Agency for Medicines and Health Products Safety (ANSM). Approval numbers: ID RCB 2014-A01953-44; ANSM reference 141587B-31.

A study involving the same participants, but examining learning mechanisms in a different task during the same experimental session, has previously been published (Di Rienzo et al., 2019). Due to the low variability observed in the behavioral measures in the BLLT, we expected low variability in neural responses, which justified using a smaller sample size, consistent with previous MEG studies on APA in the BLLT (N=15, Ng et al. (2013a)). As no prior study had specifically estimated the correlation between beta activity and inhibition strength, we based the effect size on a related study by Jana et al. (2020), which reported a correlation of r = 0.66 between beta burst timing and EMG suppression timing in a stop-signal task. Using this effect size, the power analysis indicated that a sample of 12 subjects would be sufficient to achieve 80% power at an alpha level of 0.05 for a one-sided correlation.

### Experimental setting and task

The experimental setup of the BLLT has been described in previous studies (Barlaam et al., 2011; Di Rienzo et al., 2019). Participants were seated under the MEG helmet, with a wooden table placed above their right knee, allowing the right arm to rest comfortably. The left arm was positioned alongside the trunk, with the elbow supported and constrained to allow only upward and downward rotation. A non-magnetic wristband equipped with a vacuum switch system (30 kN/m²) strain gauge was worn on the wrist, enabling an 850 g load to be placed on it in the voluntary unloading condition, or suspended and released via 3.5 kN/m² air pulses in the imposed unloading condition. During both voluntary and imposed unloading conditions, participants were instructed to maintain the left forearm in a horizontal, semi-prone position, as if supporting a tray, thereby defining the reference frame to be preserved.

In the voluntary unloading condition, the participants positioned their right hand above the load to be lifted. Each trial started with a two-second fixation on a light spot projected onto the wrist to limit eye movements during lifting. Upon light extinction (fade-out), participants were instructed to lift the load placed on top of the wristband at any time, raising it up to approximately ten centimeters above the device. After a few seconds of holding the load, participants had to return it to its initial position, completing the trial. Since participants performed the unloading themselves, APA minimized forearm destabilization, resulting in low elbow rotation following unloading. In this condition, APA are characterized by anticipatory EMG inhibition in the forearm flexor muscles (*Biceps brachii* and *Brachioradialis*), which in adults typically occurs within 50 ms before unloading (Hugon et al., 1982; Viallet et al., 1987; Schmitz et al., 2002; Barlaam et al., 2012).

In the imposed (control) unloading condition, the experimenter released the load at a random time after light extinction, preventing participants from anticipating the release and thus minimizing the engagement of APA. In contrast to voluntary unloading, imposed unloading results in a significant passive upward rotation of the elbow and inhibition of the forearm flexor muscles after unloading onset, known as the unloading reflex (Hugon et al., 1982; Schmitz et al., 2002), which serves to stabilize posture after disturbance.

During the experimental session, participants completed three conditions in the following order: imposed unloading, voluntary unloading, and a learning condition previously analyzed in Di Rienzo et al. (2019). For the purposes of the present study, only the imposed and voluntary unloading conditions were examined. The voluntary condition was preceded by a short training session to minimize adaptation and ensure stable performance. In both voluntary and imposed conditions, 90 trials were organized into blocks of ten, with approximately one minute of rest between blocks to reduce fatigue.

### Behavioural data acquisition

#### Detection of load-lifting timing

The onset of load-lifting (unloading, zero reference time point) was automatically identified as the initial deflection in the force signal recorded by the force plate sensor attached to the non-magnetic wristband, using a threshold function in CTF® DataEditor software. The automatically assigned label was then visually inspected and manually corrected if needed.

#### Elbow-joint rotation

Upward rotation of the elbow joint following load release or lifting was measured across all trials and conditions using a copper potentiometer aligned with the elbow joint axis and sampled at 20 kHz. The signal was resampled at 600 Hz for analysis. The deflection amplitude was recorded in arbitrary units provided by the equipment.

#### Electromyography

Electromyography (EMG) data were collected at a sampling frequency of 600 Hz using passive, MEG-compatible Ag/AgCl bipolar surface electrodes (2.5 mm² surface area) positioned over the left *Brachioradialis*, *Biceps brachii*, and *Triceps brachii* on the postural side, as well as the right arm *Biceps brachii*. EMG recordings were obtained following skin preparation and electrode placement procedures consistent with previously published recommendations for surface EMG sensors and sensor placement procedures (Hermens et al., 2000). The electrode sites were cleaned with alcohol wipes and thoroughly dried to reduce impedance. Bipolar Ag/AgCl surface electrodes were placed over the muscle belly, aligned with muscle fibers, with an inter-electrode distance of ∼20 mm (center-to-center). Care was taken to avoid tendon regions and adjacent muscles to minimize cross-talk. A reference electrode was placed over a bony prominence. Electrode placement was guided by anatomical landmarks and confirmed by palpation during specific isometric contractions. For the *Biceps brachii*, electrodes were positioned on the anterior upper arm (∼50% of the distance between the acromion and antecubital fossa) and confirmed during resisted elbow flexion with forearm supination. For the *Brachioradialis*, electrodes were placed on the lateral forearm (∼one-third of the distance between the lateral epicondyle and radial styloid) and confirmed during resisted elbow flexion with the forearm in a neutral position. For the *Triceps brachii*, electrodes were positioned on the posterior upper arm (∼50% of the distance between the acromion and olecranon, over the long head) and confirmed during resisted elbow extension. Standard skin preparation procedures were applied to reduce impedance and ensure adequate signal quality, consistent with common practice in surface EMG studies. Impedance was controlled (<10 kΩ) using an external impedance meter (GRASS). EMG signals were transmitted via shielded cables to a differential amplifier outside the magnetically shielded room and hardware-filtered at 0.16 Hz. Common-mode noise was reduced through differential amplification. EMG signals were synchronized with the MEG signal. Visual inspection during rest periods confirmed an adequate signal-to-noise ratio.

To investigate anticipatory muscle inhibition in this study, we analyzed EMG data from the left *Biceps brachii*, a postural elbow flexor that suppresses activity in anticipation of unloading in the BLLT, as indicated by decreased EMG activity (Massion et al., 1999). Though the left *Brachioradialis* exhibits similar dynamics, being inhibited in the same trials as the *Biceps brachii,* with comparable onset and duration (Barlaam et al., 2011), we focused our analysis on the *Biceps brachii* due to its higher signal-to-noise ratio. Considering their closely matched dynamics, we however suggest that inhibition of both elbow flexors, *Biceps brachii* and *Brachioradialis*, is governed by similar postural control mechanisms.

### Behavioural data preprocessing

#### Elbow rotation

To evaluate postural stabilization as an indicator of APA efficacy in the voluntary unloading condition, we introduced two metrics: Peak Elbow Rotation and Elbow Rotation Decline, representing the two principal directions of forearm destabilization (see Results, section *Behavioural data*). We assumed that in trials when APA are effective, both Peak Elbow Rotation and Elbow Rotation Decline would exhibit values close to zero, indicating optimal postural stabilization.

To estimate Peak Elbow Rotation and Elbow Rotation Decline in the voluntary unloading condition, the time point of maximal elbow deflection was visually identified in the time series using the ‘annotations’ tool in MNE-Python. This step was performed manually because, in most trials, peak elbow rotation was too small to be reliably detected with automatic algorithms. To minimize bias in peak annotation, the procedure was preceded by a training session with a specialist experienced in this type of data. Before annotating the peaks, trials from each participant were previewed to assess within-subject variability. The peak was detected within 500 ms after unloading onset and was typically present shortly after the end of unloading. After initial detection, peaks were visually re-inspected on a separate day, with participants randomized for each review to minimize bias. All ambiguous trials were subsequently examined with a specialist experienced in this type of data. Trials containing artifacts that prevented accurate estimation of the peak or baseline used for elbow rotation measures calculation were excluded.

The time series were then epoched from -1.7 to 1.2 s relative to the annotated peak of elbow deflection. Peak Elbow Rotation was computed as the average of baseline-corrected values around the point of maximal deflection. Elbow Rotation Decline was computed automatically as the average of baseline-corrected minimum values preceding the maximal elbow deflection (see Supplementary Materials for details on the time intervals used).

Elbow Rotation Decline and Peak Elbow Rotation were calculated individually for each trial, followed by visual inspection of the time intervals used for their calculation (see examples in Supplementary Fig. S1). For demonstration purposes, Figure 1B shows the resulting amplitudes of Peak Elbow Rotation and Elbow Rotation Decline in three example trials. Trials containing severe artifacts that precluded reliable estimation of elbow rotation measures were identified and excluded. The number of such trials per participant ranged from 0 to 5 (mean = 1.25, SD = 1.39). All analyses involving elbow rotation data were conducted on the remaining artifact-free trials.

**Figure 1.**
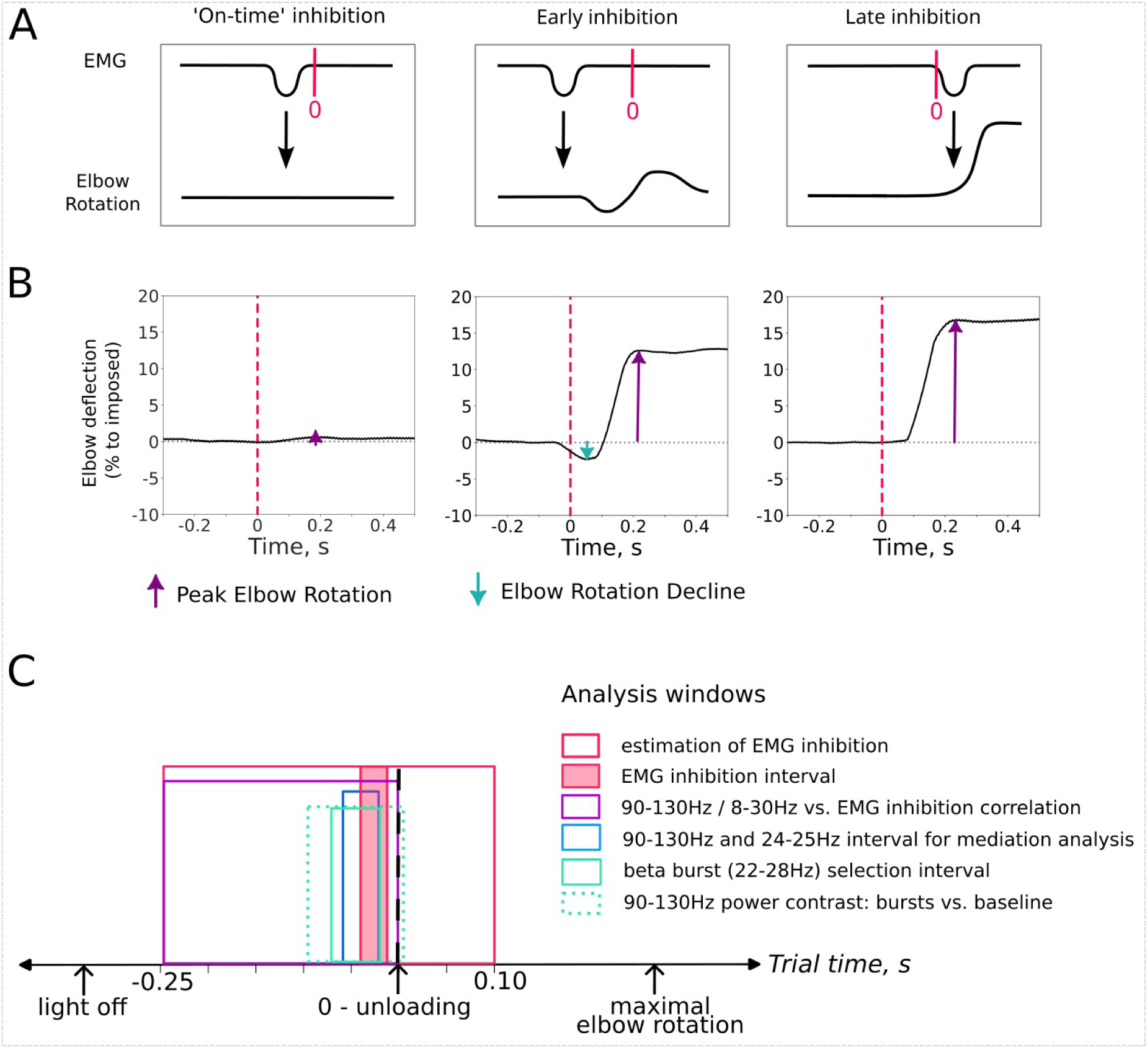
Estimation of elbow deflection parameters and their predicted link to the timing of *Biceps brachii* inhibition, with a schematic showing the intervals used in the main study analyses. A: Schematic illustration of three primary possible scenarios for *Biceps brachii* inhibition timing and its hypothetical impact on elbow rotation. See Methods (*Estimation of the EMG inhibition* section) for details. **B:** Single-trial examples aligned to unloading onset (time = 0), schematically illustrating the amplitudes of Peak Elbow Rotation and Elbow Rotation Decline. Values are presented as percentages of elbow deflection during voluntary unloading relative to the individual maximum, estimated as the median peak elbow rotation in the imposed condition. **C.** Schematic illustration of the time intervals used in the main analyses and for estimating key measures during voluntary unloading. Exact intervals and their justification are provided in the Methods.

In the imposed condition, to assess the postural destabilization triggered by the experimenter’s unexpected release of the load, only upward elbow rotation was measured, as a decrease in elbow rotation was not observed in this condition. Peak Elbow Rotation was defined as in the voluntary condition, except that the prominent peak allowed fully automatic detection by taking the maximal elbow deflection within the trial window. This approach was reliable since each trial contained a single peak that was consistently and substantially above baseline.

#### Electromyography (EMG)

For this study, we analysed EMG recordings from the left *Biceps brachii*. Preprocessing followed established protocols (Schmitz et al., 2002; Barlaam et al., 2011, 2018) and included epoching from -1 to 0.5 s relative to the onset of unloading, mean subtraction, and band-pass filtering using a finite impulse response (FIR) filter with a Hamming window in the 25-150 Hz range, chosen after inspecting group-averaged muscle power spectral density. The filtered signals were then rectified to enable visualization of *Biceps brachii* inhibition as a reduction in EMG amplitude (see Fig. 2A, right panel). The signal was log10-transformed and baseline-corrected (-1 to -0.5 s). To determine whether EMG significantly decreased at the group level, we performed a one-tailed permutation cluster test comparing the resulting EMG amplitude to zero in the -0.25 to 0.1 s and 0 to 0.35 s windows around unloading for the voluntary and imposed conditions, respectively.

**Figure 2.**
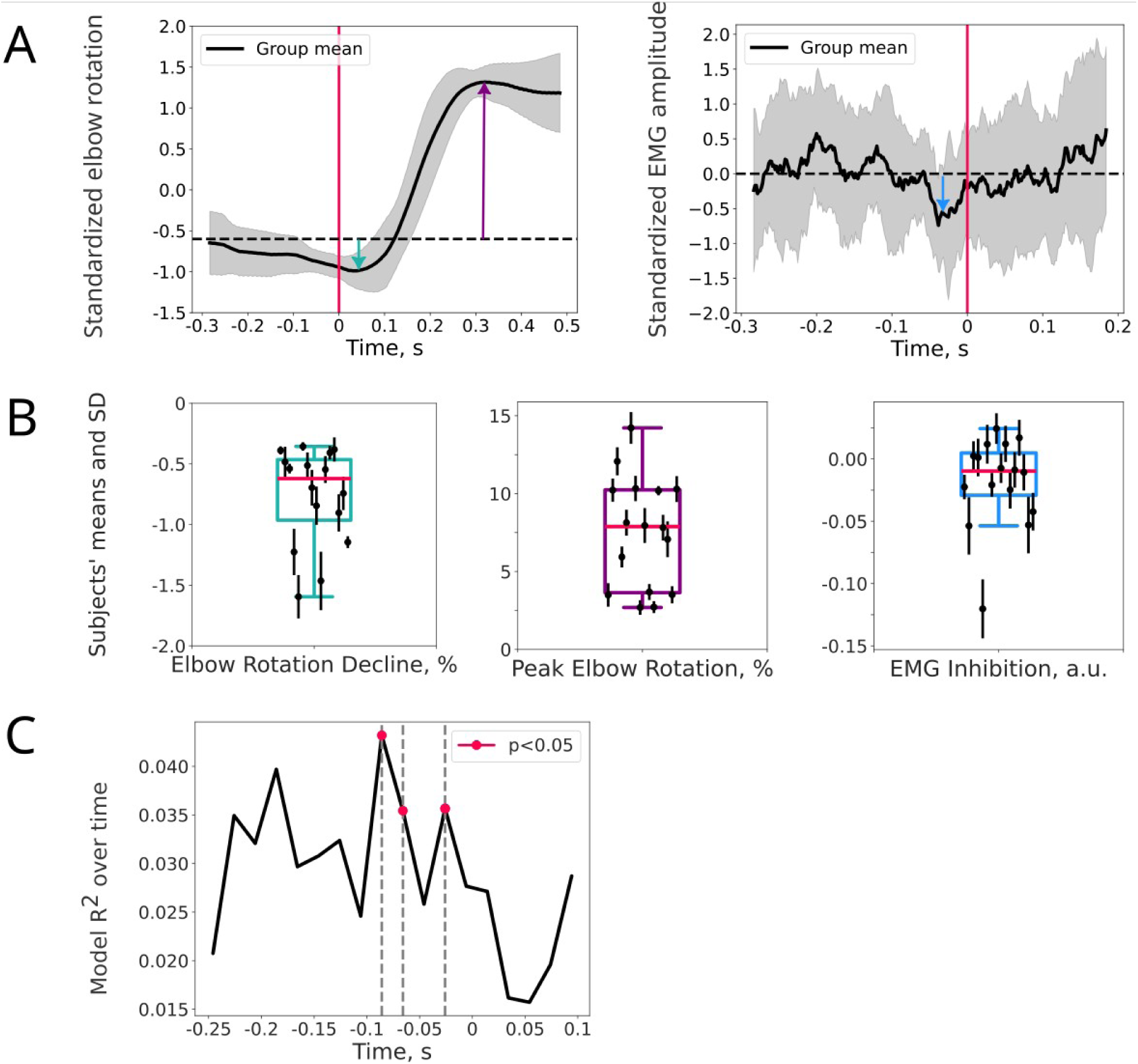
Modulation of EMG and elbow rotation amplitudes over time, with the distribution of derived parameters across subjects, and a model showing intervals where EMG is predicted from elbow rotation. Data in panels **A** and **C** are aligned to unloading onset (time = 0). **A:** Standardized elbow rotation and *Biceps brachii* EMG. Group means (black line) and SD (gray shading) are shown, with colored arrows indicating the amplitudes of behavioural measures: Elbow Rotation Decline (green), Peak Elbow Rotation (purple), and EMG inhibition (blue). **B:** Participant-wise mean (dots) and SD (whiskers) values for each measure, with boxplots indicating between-subject variability. The red line marks the median, box edges show quartiles (Q1 and Q3), and whiskers extend to 1.5 times the interquartile range. **C:** Group-averaged R² values from linear regression models fitted in 30-ms sliding time windows (dependent variable: EMG amplitude; independent variables: Peak Elbow Rotation, Elbow Rotation Decline). Red markers and gray dashed lines denote time points (time-averaged 30-ms intervals) at which the relationship between EMG amplitude and elbow rotation parameters reached significance.

To identify the time interval during which *Biceps brachii* inhibition was optimal for forearm stabilization in the voluntary condition, the EMG signal was further smoothed using a 30-ms moving average with 33% overlap to improve the reliability of the inhibition estimates and predictions. The 30-ms window was guided by previous evidence indicating that *Biceps brachii* inhibition typically lasts ∼50-100 ms in adults (Barlaam et al., 2011; Schmitz et al., 2002). This window thus captured a shorter interval than the inhibition event and was used to balance temporal resolution and the robustness of the estimates, thereby enabling the identification of a reliable interval during which inhibition was associated with optimal postural stabilization. The use of a 33% overlap allowed us to test neighboring intervals and refine identification of the optimal interval.

#### Estimation of the EMG inhibition

Since high noise levels in the EMG signal prevented accurate trial-by-trial assessment of muscle inhibition, we instead aimed to quantify inhibition strength during the period when it most effectively contributed to subsequent postural stabilization – specifically, when inhibition was followed by the maintenance of a stable horizontal forearm position. This involved identifying the time interval in which *Biceps brachii* inhibition had the strongest effect on both Elbow Rotation Decline and Peak Elbow Rotation, bringing both values close to zero and indicating a stabilized posture.

The anticipatory muscle inhibition observed in this paradigm likely reflects the suppression, or adjustment, of the ongoing postural command responsible for elbow flexor control during load support, finely tuned to maintain the forearm in a horizontal position despite the loss of opposing force due to load release. Its effectiveness depends critically on timing, which must be optimal: not too early, to ensure forearm stability until the load is released, and not too late, to prevent destabilization caused by the sudden change in force. Figure 1A provides a schematic overview of these three timing scenarios: early, on-time, and late *Biceps brachii* inhibition, along with their expected behavioural effects.

To investigate the mechanisms underlying the most efficient strategy, i.e., on-time inhibition, which is predominant in adult participants (Schmitz et al., 2002), we aimed to determine the time at which muscle inhibition results in the smallest negative elbow deflection (Elbow Rotation Decline) and lowest Peak Elbow Rotation.

To this end, we fitted linear regression models for each participant across trials, using Elbow Rotation Decline and Peak Elbow Rotation as predictors of *Biceps brachii* EMG. The OLS() function from the statsmodels.regression.linear_model module (statsmodels library, v. 0.14.1) was used to fit a separate linear regression model at each time point of the smoothed EMG signal (see above). The F-value was used as the evaluation metric for each regression model, and the coefficient of determination (R²) was used to assess the quality of the prediction. F-values were log10-transformed to normalize their distribution and baseline-corrected (-1 to -0.5 s). Therefore, positive baseline-corrected F-values indicated time intervals where Elbow Rotation Decline and Peak Elbow Rotation predicted *Biceps brachii* EMG more strongly than its baseline activity. A one-sample permutation cluster test was used to assess whether the positive, baseline-corrected F-values significantly differed from zero across subjects, within the time interval of -0.25 to 0.1 s relative to unloading.

In the imposed condition, in which the timing of load release could not be anticipated, the unloading reflex is elicited, characterized by inhibition of the *Biceps brachii* (Hugon et al., 1982; Schmitz et al., 2002), which serves to stabilize posture following the disturbance. To quantify the strength of *Biceps brachii* inhibition in this condition, EMG amplitude was averaged over the time window showing a significant group-level decrease, thereby providing an estimate of reactive EMG inhibition strength across trials and participants.

### Neuroimaging data acquisition and analysis

Participants underwent both magnetoencephalography (MEG) and structural magnetic resonance imaging (MRI). Structural MRI data were acquired to enable MEG-MRI coregistration and subsequent source localization. Accordingly, MRI acquisition and preprocessing are described first, followed by the MEG acquisition and analysis procedures.

#### MRI data acquisition and preprocessing

Structural MRI (voxel size 0.9 mm x 0.9 mm x 0.9 mm; TR=3500 ms, TE=2.24 ms) data were obtained from all participants using a 3T Siemens Magnetom scanner (CERMEP, France - MAGNETOM Prisma, Siemens HealthCare). T1-weighted images were preprocessed with the ‘recon-all’ procedure from FreeSurfer software (version 6.0.0; Fischl et al., 2002), which includes motion correction, intensity normalization, removal of non-brain tissue, cortical surface reconstruction, and subcortical segmentation.

The cortical surface was parcellated into 450 regions using the ‘HCPMMP1’ atlas (Glasser et al., 2016), as implemented in MNE-Python software (v. 1.7.0; Gramfort et al., 2013). Source-level MEG analysis was then restricted to brain regions previously associated with voluntary unloading in the BLLT task (Schmitz et al., 2005; Ng et al., 2011, 2013a, 2013b). To investigate anticipatory inhibition mechanisms, our analysis specifically targeted the right-hemisphere network, contralateral to the load-supporting arm, where inhibition occurs. Although anticipatory inhibition is inherently linked to bimanual coordination in the BLLT, focusing on the right-hemispheric network allowed us to investigate the efferent pathways involved in postural *Biceps brachii* inhibition and APA, which primarily rely on the hemisphere contralateral to the postural forearm (Massion, 1992; Viallet et al., 1992). A full list of the regions of interest is provided in Supplementary Table S1.

To examine the potential contribution of subcortical regions previously implicated in the BLLT (Viallet et al., 1989; Schmitz et al., 2005; Ng et al., 2011; 2013a; 2013b) to anticipatory *Biceps brachii* inhibition, the basal ganglia and cerebellum were included in MEG source-level analysis, together with cortical regions. These regions were automatically parcellated using the Aseg atlas in FreeSurfer which included the cerebellar cortex and basal ganglia (caudate nucleus, putamen, and globus pallidus).

#### MEG data acquisition

Magnetoencephalography (MEG) recordings were acquired using a CTF-MEG system (CERMEP, France - CTF MEG Neuro Innovations, Inc), equipped with 275 radial gradiometers positioned to cover the scalp, along with 29 reference channels for correcting ambient noise. The MEG signals were digitized at a 600 Hz sampling rate and low-pass filtered between 0 and 150 Hz. Head position was tracked continuously using three coils placed on the nasion and preauricular points before recording.

### MEG sensor-level data analysis

#### Preprocessing steps

MEG data preprocessing and analysis were performed using MNE-python software (v. 1.7.0; Gramfort et al., 2013). Raw MEG data from the voluntary unloading condition (three sessions per participant), aligned to the initial head position within each session, were concatenated. Visual inspection confirmed that head movements did not exceed 1 cm along the x, y, or z axes. A detailed analysis further showed that head position remained stable across all three coordinates during the anticipatory period (-0.5 to 0 s relative to unloading), which was used in subsequent analyses.

The data were high-pass filtered at 1 Hz before applying Independent Component Analysis (ICA) using the ICA() function (number of components: 70, method: ’picard’, maximum number of iterations: 1000, reference channels were included in the IC estimation). Components reflecting biological artifacts (blinks, heartbeat, muscle activity) were identified and excluded based on visual inspection of their time courses and topographies. To detect components contaminated by intermittent environmental noise, the find_bads_ref() function (threshold=1.5) was applied to the MEG reference channels. Identified components were then removed from the unfiltered copy of the raw data. The average number of excluded components (mean ± SD) was as follows: EOG-related: 2.06 ± 0.24; ECG-related: 1.25 ± 0.56; EMG-related: 1.44 ± 1.54; artifact noise: 6.56 ± 3.46.

The raw data were epoched from -1.7 to 1.2 s relative to unloading onset. Note that, in the MEG data, no baseline correction was applied at this stage or in any subsequent analyses. Only trials in which the unloading onset was detectable and the elbow rotation signal did not exhibit severe artifacts were retained for further analysis (number of dropped trials per subject, mean ± SD: 1.19 ± 1.18). Data of 275 axial gradiometers were selected for the analysis. Epochs were visually inspected and those contaminated by instrumental noise and myogenic artifacts were excluded (number of dropped trials per subject, mean ± SD: 2.00 ± 2.89). Several trials were additionally excluded from the elbow rotation analyses due to the presence of artifacts in time intervals critical for reliably estimating Peak Elbow Rotation and/or Elbow Rotation Decline (see above). The final average number of trials per participant was 86.94 ± 2.97 (mean ± SD).

For the imposed (control) condition, preprocessing included the same steps, with epoching performed in the time interval of -0.5 to 0.5 s relative to unloading onset, resulting in a total of 87.00 ± 5.55 (mean ± SD) trials per subject.

### MEG source-level data analysis

#### Individual brain models and inverse solution

To align MEG sensors and head coordinates with each participant’s MRI in a common coordinate system, we used the coregistration procedure implemented in mne.gui.coregistration(). Three HPI coil locations (nasion, left, and right) were directly marked on the participant’s skin prior to MEG acquisition, and corresponding MRI-visible aquagel markers (3 mm) were placed at the same positions before MRI. These markers were clearly visible on the individual MRI head surface, allowing for accurate coregistration.

A single-layer boundary element model (BEM) was created with 20,484 vertices across both hemispheres. A mixed source space was defined, combining a surface-based source space with ’ico4’ spacing yielding 5,124 cortical vertices in both hemispheres and a volume source space with 5 mm spacing, which included the cerebellum and basal ganglia comprising a total of 1,300 sources. The forward solution was computed for sources with a minimal distance of 5 mm from the inner skull surface.

Notch filters at 50 and 100 Hz were applied to the raw data after ICA to reduce powerline noise. The data were then band-pass filtered at 25-150 Hz for gamma activity analysis, 1-90 Hz for alpha-beta power analysis, and 1-150 Hz for alpha-beta vs. gamma correlation using zero-phase FIR filters with a Hamming window. The filter length was set to ‘auto’ with the ‘firwin’ design and ‘reflect_limited’ padding. The resulting data were epoched as described in the preprocessing step, and bad trials were excluded based on prior annotations.

The data rank was computed from the epoched data, with a singular value tolerance of 1e-6 applied relative to the largest singular value. The data covariance matrix was then calculated using this rank with the ‘empirical’ method for the time interval from -1 to 0.1 s relative to unloading onset to capture the brain response associated with anticipatory motor control. Since only gradiometer channels were used and the beamformer approach was adopted, noise covariance estimation for source reconstruction was not necessary. A unit-gain Linearly Constrained Minimum Variance (LCMV) beamformer spatial filter was computed to optimize power orientation, with a regularization coefficient of 0.05 and enabling the rank reduction. The resulting filter was applied to each epoch individually.

#### Time-frequency analysis

Time-frequency analysis was performed over the full time window from -1.7 to 1.2 s, with symmetric padding of 0.5 s added on both sides of the time array. The resulting data were then cropped to the window of interest (-0.5 to 0.25 s) relative to unloading.

For gamma power estimation, the Multitaper method (using the tfr_array_multitaper() function) was applied to the time series of individual-trial source activity reconstructed with the LCMV beamformer. The analysis was conducted in the 90-130 Hz frequency range, using 7 cycles, a time_bandwidth parameter of 4, and a frequency resolution of 1 Hz. The resulting power estimates were log10-transformed and averaged across the frequency dimension to yield a mean high-gamma power estimate for the 90-130 Hz band.

For power estimation in the alpha-beta range, the adaptive Superlets algorithm was used (https://github.com/irhum/superlets; Moca et al., 2021). For each trial and brain source, power was estimated in the 1-80 Hz frequency range (base cycle = 3, orders 1-20). Periodic power was then extracted per trial and source using the Specparam tool (‘FOOOF’; Donoghue et al., 2020) as the difference between total power and the aperiodic fit, derived from trial-averaged spectra in 2-80 Hz, which was selected based on visual inspection of spectral densities. For further analysis, the 8-30 Hz alpha-beta band was extracted.

#### Correlational analyses

Previous studies suggest that anticipatory elbow flexor inhibition during voluntary unloading may be attributed to reduced excitability, rather than enhanced inhibitory drive, in the contralateral brain region governing muscle control (Kazennikov et al., 2005, 2006). To test this, we used Spearman’s rank correlation to relate our estimate of anticipatory inhibition – EMG inhibition (see Results, *Identification of the optimal timing of anticipatory EMG inhibition* section) – to high-gamma power over time, averaged in the 90-130 Hz band and used as a proxy for cortical excitability. High-gamma activity – typically defined as neural activity within a broad frequency range above 60 or 80 Hz – has been shown to correlate closely with neuronal spiking in both sensory (Ray et al., 2008; Ray and Maunsell, 2011; Suffczynski et al., 2014) and motor networks (Yazdan-Shahmorad et al., 2013), supporting its use as an indicator of regional excitability. To assess the link between muscle inhibition and the proposed index of excitability, correlations between EMG inhibition and high-gamma power were computed at each time point between -0.25 and 0 s relative to unloading and for each cortical source within the right anticipatory motor network (see Supplementary Table S1), to capture activity associated with anticipatory EMG inhibition. A one-tailed, one-sample permutation cluster test was used to assess whether stronger EMG inhibition is linked to reduced high-gamma power in the selected network.

In the imposed condition, the correlation between high-gamma power and reactive EMG inhibition was evaluated in the 0-150 ms interval after unloading. This interval was selected to capture activity associated with postural *Biceps brachii* inhibition in this condition, representing an unloading reflex elicited by forearm destabilization, peaking around 100 ms after unloading (Hugon et al., 1982).

To investigate the potential involvement of additional brain regions in the regulation of anticipatory muscle inhibition mediated by alpha- and beta-band activity, we computed Spearman’s rank correlations between periodic power and EMG inhibition at each source vertex within the right anticipatory motor network, across time-frequency points in the 8-30 Hz range and the -0.25 to 0 s time window. To further assess the contribution of subcortical structures, the same correlation analysis was performed for the right cerebellum and basal ganglia. Separate one-tailed, one-sample permutation cluster tests were conducted to determine whether stronger EMG inhibition was associated with increased alpha- and beta-band power across both the right anticipatory motor network and the subcortical structures.

Since stronger EMG inhibition correlated with reduced high-gamma power in the medial SMA (see Results), we aimed to specifically test whether inhibitory effects in the SMA were mediated by alpha- or beta-band activity. To assess whether stronger muscle inhibition is related to alpha- or beta-band power, we estimated Spearman correlations between periodic power and EMG inhibition across time-frequency points in the 8-30 Hz range and the -0.25 to 0 s interval at each of the 10 vertices within the SMA cluster label. To evaluate whether modulation of SMA excitability, as reflected in high-gamma activity, is mediated by alpha- or beta-band activity, we further estimated correlations between total alpha-beta (8-30 Hz) and high-gamma (90-130 Hz) power within the selected SMA cluster. To control for shared variance associated with broadband power fluctuations, we computed partial Spearman’s correlation coefficients at each vertex using the partial_corr() function from Pingouin (v. 0.5.3), with mean total power in the 1-130 Hz range included as a covariate (Spaak et al., 2012). The resulting correlation coefficients were then averaged across cluster vertices. One-tailed, one-sample permutation cluster tests were conducted to determine whether stronger alpha-beta power was associated with greater EMG inhibition and reduced high-gamma power in the SMA.

#### Mediation analysis

To further clarify the potential mechanism underlying the observed interplay between stronger EMG inhibition, reduced high-gamma power, and greater high-beta power in the SMA (see Results), we proposed a theoretical model in which high-beta activity underlies the reduction of high-gamma activity in the SMA, which in turn results in muscle inhibition, thereby accounting for the observed relationships between greater beta activity and both reduced high-gamma and stronger EMG inhibition. This model therefore suggests that beta activity is associated with EMG inhibition indirectly, by exerting inhibitory control over the SMA, as reflected in high-gamma suppression, consistent with prior reports of an inverse relationship between beta and high-gamma/spiking activity in sensorimotor areas (Ray et al., 2008; Lundqvist et al., 2016; Riehle et al., 2018). In turn, the model implies that the reduction in high-gamma activity, indexing decreased SMA excitability, relates to the strength of anticipatory *Biceps brachii* inhibition, thereby highlighting a potential role for the SMA in controlling this muscle.

To test the proposed model, we conducted a mediation analysis. Specifically, we tested whether high-gamma power (mediator, M) mediated the relationship between high-beta power (independent variable, X) and EMG inhibition (dependent variable, Y). The analysis focused on the ‘peak vertex’ within the SMA cluster region, defined as the vertex showing the strongest correlation between high-gamma power and EMG inhibition.

For mediation analysis, we selected beta power in the 24-25 Hz range from a broader 18-25 Hz cluster showing a correlation with EMG inhibition (Fig. 3B). This narrower range was selected as it overlapped with the high-gamma correlation effect (Fig. 3A) and the EMG inhibition calculation interval (-26±15 ms relative to unloading). This suggested that the relationship between beta activity in this frequency range and EMG inhibition could be partially explained by the negative correlation between beta and high-gamma power.

**Figure 3.**
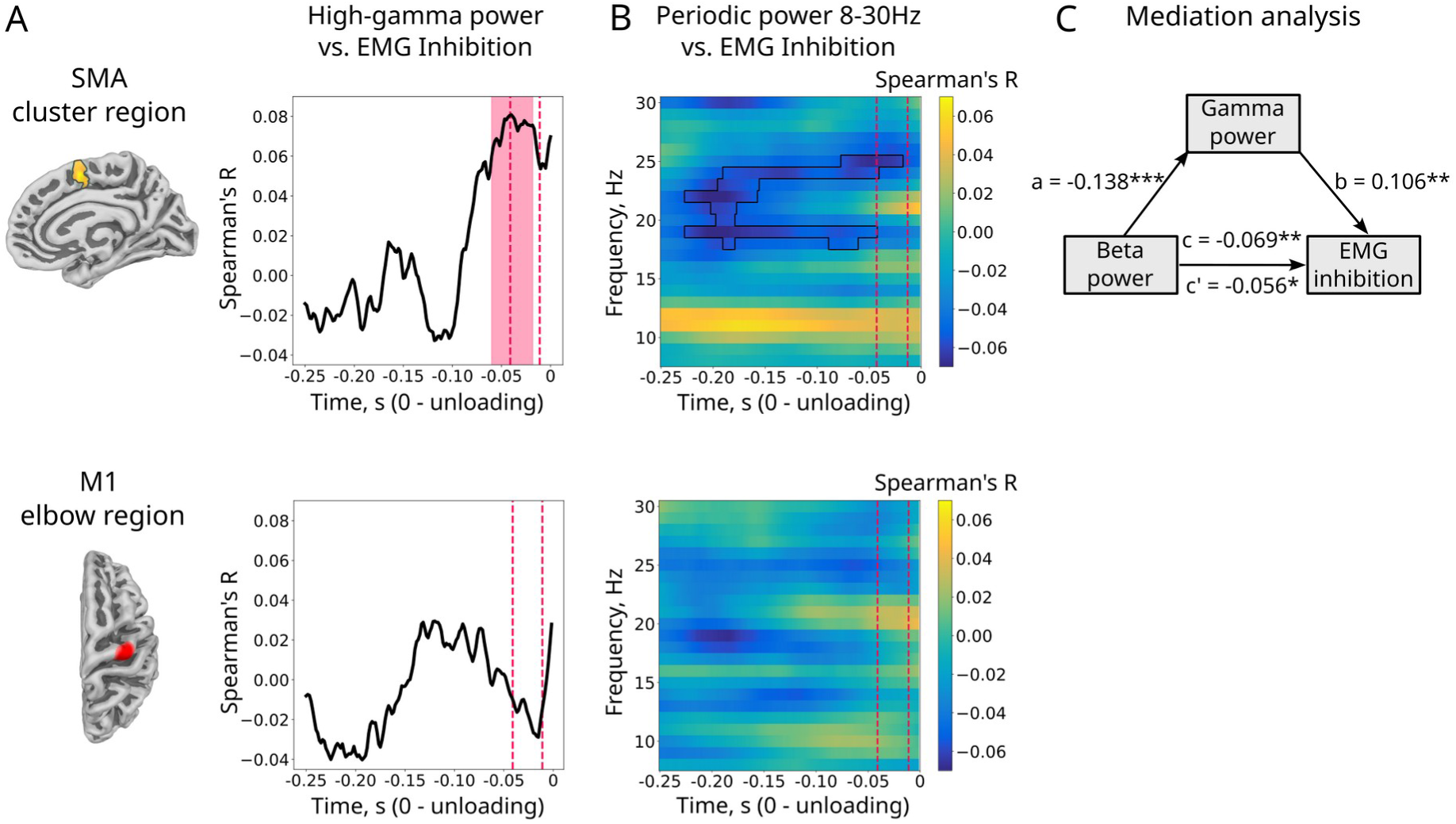
EMG inhibition correlates with high-gamma and beta power in the SMA, with high-gamma power mediating the relationship between high-beta power and EMG inhibition. Data in panels **A** and **B** are aligned to unloading onset (time = 0). A: *Upper panel:* A cluster-level significant correlation effect between high-gamma power (90-130 Hz) and EMG inhibition (averaged over a -26±15 ms interval, indicated by red dashed lines) with the strongest effect observed in the medial SMA region (MNI305 peak: [8, 3, 52]). The thick black line represents the correlation time course within this cluster, with the red-shaded area highlighting its time span (p < 0.05, corrected). *Lower panel*: The right M1 elbow area (MNI305 peak: [32, -20, 66]) and its correlation time course, which did not reach significance. **B:** Spearman’s correlation coefficients (R) between 8-30 Hz periodic power and EMG inhibition in the SMA cluster (upper) and M1 elbow region (lower). Thin black lines indicate significant correlation effect (p < 0.05, corrected). **C:** Mediation analysis results showing the relationships between Beta power (independent variable), High-gamma (90-130 Hz) power (mediator), and EMG inhibition (dependent variable). Linear regression coefficients with the respective p-values are reported. **\*p<0.05; **p<0.01; ***p<0.001**

Total (without aperiodic component subtraction) beta and high-gamma power were averaged over their respective frequency bands – 24-25 Hz for beta and 90-130 Hz for gamma – and across the time interval in which a significant correlation between gamma power and EMG inhibition was observed (-0.06 to -0.018 s relative to unloading). Notably, this time window overlapped with the interval during which beta power (24-25 Hz) was correlated with EMG inhibition (see Fig. 3B). The use of a unique time interval for estimation of both the independent variable and the mediator is justified by the fast conduction velocity of neural signals (less than 1 ms), which is below the time resolution of the current data, making it impossible to differentiate their exact timings. Before fitting the models, beta power was adjusted by regressing out (LinearRegression() function, ‘sklearn’, v. 1.4.2) the total power averaged across a broad frequency range (1-130 Hz) to mimic the partial correlation approach used for the beta power vs. gamma power correlation (see above). The resulting beta and gamma powers were further scaled across subjects to yield standardized mediation parameters.

The mediation analysis employed bootstrapping with 10,000 permutations to obtain robust estimates of the mediation effects. To account for random effects, generalized additive models (GAMs) with linear fixed effects (gamma power, beta power) and random effects per subject (random intercepts, random gamma and beta slopes) were fitted using restricted maximum likelihood (REML).

To examine whether high-gamma power mediates the relationship between high-beta power and EMG inhibition, we conducted a mediation analysis using four models summarized in Table 1. This approach allowed us to assess whether the association of high-beta power with EMG inhibition may be partly explained by its association with high-gamma power reduction.

**Table 1.** Mediation analysis models.

| Model | Relationship | Formula (mgcv) |
| --- | --- | --- |
| A | $X \rightarrow M$ | $\text{Gamma} \sim \text{Beta} + s(\text{Subject}, \text{bs} = 're') + s(\text{Subject}, \text{Beta}, \text{bs} = 're')$ |
| C' | $X + M \rightarrow Y$ | $\text{Inhibition} \sim \text{Beta} + \text{Gamma} + s(\text{Subject}, \text{bs} = 're') + s(\text{Subject}, \text{Beta}, \text{bs} = 're') + s(\text{Subject}, \text{Gamma}, \text{bs} = 're')$ |
| B | $M \rightarrow Y$ | $\text{Inhibition} \sim \text{Gamma} + s(\text{Subject}, \text{bs} = 're') + s(\text{Subject}, \text{Gamma}, \text{bs} = 're')$ |
| C | $X \rightarrow Y$ | $\text{Inhibition} \sim \text{Beta} + s(\text{Subject}, \text{bs} = 're') + s(\text{Subject}, \text{Beta}, \text{bs} = 're')$ |

*Inhibition*: EMG inhibition; *Gamma*: gamma power averaged over 90-130 Hz; *Beta*: adjusted beta power averaged over 24-25 Hz; $s(\text{Subject}, \text{bs} = 're')$ : subject-based random intercept; $s(\text{Subject}, \text{Beta}, \text{bs} = 're')$ : subject-specific random slope for beta power; $s(\text{Subject}, \text{Gamma}, \text{bs} = 're')$ : subject-specific random slope for gamma power.

In short, model A examined the relationship between high-beta power (X, independent variable) and high-gamma power (M, mediator). Model B examined the relationship between the mediator (high-gamma power) and EMG inhibition (dependent variable) without including the independent variable. Model C examined the relationship between the independent variable (high-beta power) and EMG inhibition without including the mediator. Model C’ examined the combined relationship of the independent variable (high-beta power) and the mediator (high-gamma power) with EMG inhibition, allowing estimation of the portion of the association between high-beta power and EMG inhibition that can be accounted for by high-gamma power. Details justifying the linearity assumptions of these models are provided in the Supplementary Methods.

#### Beta bursts extraction

To extract beta bursts at the vertices of interest, we employed the burst detection pipeline developed by Szul et al. (2023; https://github.com/danclab/burst_detection). Time-frequency decomposition and aperiodic component extraction were performed using Superlets and Specparam, respectively, as described above (see Methods, *Time-frequency analysis* section). Unlike previous steps, a frequency resolution of 0.25 Hz was employed in the Superlet analysis to improve burst detection accuracy. To avoid edge effects, the algorithm was applied to the 10-33 Hz range. Beta bursts were then selected within the 13-30 Hz frequency band within a -0.8 to 0.4 s window, relative to unloading. To verify the accuracy of burst extraction, trial-wise time-frequency representations with identified bursts were visually inspected.

Bursts were extracted for each trial from the ‘peak vertex’ within the SMA cluster region, identified as showing the strongest correlation between gamma power and EMG inhibition. The choice to use the peak vertex rather than all vertices of the label was motivated by the difficulty of aggregating bursts, as some may be redundantly detected across nearby sources due to signal spread. In contrast, extracting bursts from signals averaged across all SMA vertices could introduce error, since this would require averaging raw signals and aperiodic components used by the algorithm across vertices. Thus, the peak vertex was used as a trade-off, likely capturing transient activity of the label rather than a single source due to signal spread. For this vertex, only bursts with a burst center frequency between 22-28 Hz and a burst center time between -0.077 and -0.018 s were considered for analysis. These time and frequency intervals were defined based on the observed correlation between beta power and EMG inhibition (Fig. 3B): the time window captured its temporal effect around inhibition, and the frequency range (25 ± 3 Hz) targeted the correlation peak at 25 Hz while accounting for potential variability in burst frequency estimates (see Results, *Beta burst analysis* section, for details on range selection).

#### Beta bursts analysis

To examine whether trials containing beta bursts within the selected time and frequency range also exhibit more pronounced anticipatory *Biceps brachii* inhibition, we averaged baseline-corrected EMG amplitude time courses across these trials. A one-tailed, one-sample permutation cluster test was conducted across participants within the -0.25 to 0.05 s window relative to unloading to assess whether EMG amplitude was significantly reduced during beta burst trials, indicating stronger anticipatory inhibition.

For further analysis, trials containing beta bursts were aligned to the center time of each burst. For the control trials, an equal number of no-burst trials were selected based on the lowest beta power within the time-frequency range used for burst extraction. These control trials were then aligned to the burst center times from burst trials, ensuring time shuffling in the control condition as well.

The choice to use no-burst trials with the lowest beta power as controls was motivated by the overall low proportion of burst trials (∼10% of total trials). Although consistent with previous reports of similarly low percentages (∼15%) of beta burst trials linked to successful stopping (Jana et al., 2020), this raises the problem of selecting an equal number of no-burst trials to match burst trials. Given the link between beta power and EMG inhibition, and reports of generally optimal anticipatory inhibition in adults (Schmitz et al., 2002; Barlaam et al., 2011), we reasoned that some bursts may have gone undetected in “no-burst” trials due to methodological constraints of burst detection. To maximize the use of true no-burst trials, we therefore selected no-burst trials with lower periodic beta power in the time-frequency range used for burst detection, suggesting that bursts were less likely to occur in these trials.

To test whether the occurrence of beta bursts was linked to high-gamma power suppression, we compared high-gamma (90-130 Hz) power between trials with bursts and no-burst control trials. A one-tailed, one-sample permutation cluster test (-0.05 to 0.05 s around burst centers, 90-130 Hz) was conducted to assess whether high-gamma power decreased in burst trials compared to no-burst trials.

As a complementary analysis, we investigated the right-hemispheric network supporting anticipatory inhibition. Since conventional connectivity approaches can produce spurious results for signals with transient bursts, where brief high-amplitude activity is interspersed with longer oscillation-free periods (Lundqvist et al., 2024), connectivity was estimated on MEG signals centered on beta bursts. Since this analysis was secondary to the main objectives of the study and exploratory in nature due to the limited number of burst trials, details of the analysis and results are provided in the Supplementary Materials.

#### Statistical analysis

Mediation analysis was performed in R (version 4.5.0) using the mediate() function (‘mediation’ package, v. 4.5.0) and the bam() function to fit GAMs (‘mgcv’ package, v. 1.9.0). To check the assumptions of linear regression models, several diagnostic tests were performed. Normality of residuals was assessed with the Shapiro-Wilk test (shapiro(), scipy.stats v. 1.7.3), and autocorrelation was checked using the Durbin-Watson test (durbin_watson(), statsmodels.stats.stattools v. 0.13.5). Homoscedasticity was assessed with the Breusch-Pagan test (het_breuschpagan(), statsmodels.stats.api v. 0.13.5), and multicollinearity with the Variance Inflation Factor (variance_inflation_factor(), statsmodels.stats.outliers_influence v. 0.13.5).

To assess the relationship between neural activity and behavioural parameters at individual time, frequency, and spatial points, we calculated Spearman’s rank correlation coefficient (spearmanr() function, scipy.stats, v. 1.7.3). Spearman’s correlation was chosen due to the potential presence of non-linear relationships between the pairs of variables studied.

To perform cluster analysis, group-level statistics, or to select a common set of vertices across all subjects (e.g., the SMA cluster region or the right anticipatory motor network), timecourses or correlation coefficients computed for individual brain models were morphed to the template brain, ‘fsaverage’, using the compute_source_morph() function in MNE-Python. For ‘fsaverage’, the BEM and source spaces were created using the same parameters as those applied to the sample data.

A non-parametric permutation cluster one-sample t-test was used for all cluster-level analyses, employing either permutation_cluster_1samp_test() for time-only data or spatio_temporal_cluster_1samp_test() for analyses involving spatial or frequency dimensions as well, with 1,024 permutations used to build a null distribution for cluster-level inference. To stabilize t-statistics in the presence of low variance, the ‘hat’ variance regularization was used. Clusters were formed based on adjacency in time, space, frequency, or their combinations. One- or two-tailed tests were applied depending on the hypotheses, as specified for each test in Methods. Threshold-Free Cluster Enhancement (TFCE) was used for one- or two-dimensional data (time or time-frequency; starting value: 0, step size: 0.2). For higher-dimensional data involving spatial adjacency, a cluster-forming threshold based on p = 0.001 was used to manage computational costs. We confirmed the reliability of this approach by comparing TFCE and cluster-based thresholding in a high-gamma-to-EMG inhibition correlation, which yielded similar clusters (p < 0.05) across time and within the anticipatory motor network for both methods.

## Results

### Behavioural data

#### Postural stabilization and EMG dynamics during voluntary unloading

The group-averaged elbow rotation and EMG modulation over time during voluntary unloading, including between-subject variability, are presented in Figure 2A. The time-resolved EMG signal showed a significant decrease relative to zero across participants (t(15)_min_ = -4.43, p_min_ = 0.040), spanning 38-40 ms before unloading (Fig. S2). Distributions of Peak Elbow Rotation and Elbow Rotation Decline across trials were significantly non-normal for all participants (Shapiro-Wilk test, p < 0.05). Furthermore, for both parameters, the means and SDs were low relative to each participant’s maximal elbow rotation in the imposed condition. The average ratio of voluntary to imposed elbow deflection was: Peak Elbow Rotation, 5.46 ± 7.44%; Elbow Rotation Decline, -0.43 ± 2.04% (see Fig. 2B for individual variability). Notably, the observed 5% voluntary Peak Elbow Rotation deflection closely matches earlier findings of 8% in adults (Barlaam et al., 2012), consistent with mature anticipatory control mechanisms stabilizing forearm during voluntary load-lifting.

#### Identification of the optimal timing of anticipatory EMG inhibition

Using within-participant variability in elbow deflection across trials, we aimed to determine the timing of anticipatory inhibition in the *Biceps brachii* that would optimize forearm stabilization. We assumed that optimal elbow flexor inhibition timing would minimize deviations from zero in both Peak Elbow Rotation and Elbow Rotation Decline (‘On-time’ inhibition scenario, Fig. 1A). Optimal inhibition timing was identified by fitting linear regression models predicting *Biceps brachii* EMG from trial-level variability in Peak Elbow Rotation and Elbow Rotation Decline (see Methods, *Estimation of the EMG inhibition* section). Verification of the linearity assumptions for these models is provided in the Supplementary Materials. A permutation cluster test identified significant differences in baseline-corrected F-values, with the strongest effect at three time points: -0.086 s (t(15) = 4.04, p = 0.002), -0.066 s (t(15) = 2.64, p = 0.019), and -0.026 s (t(15) = 3.9, p = 0.022) relative to unloading, indicating periods when EMG modulation was most strongly associated with elbow rotation. Group-averaged R² values for these models are shown in Figure 2C, highlighting these three key intervals. A closer examination of the correlation coefficients between EMG amplitude and the two elbow rotation metrics (see Supplementary Results, Fig. S3) revealed that only around -0.026 s was a stronger EMG suppression associated with both reduced (less negative) Elbow Rotation Decline and lower (less positive) Peak Elbow Rotation, indicative of a more stabilized forearm. The time interval 26 ± 15 ms before unloading was therefore consistent with the ‘on-time’ inhibition scenario (Fig. 1A) and corresponded to a clear reduction in group-averaged EMG activity relative to baseline (Fig. 2A, right panel). By contrast, earlier intervals (86± 15 ms and 66± 15 ms before unloading showed positive associations between EMG and Elbow Rotation Decline, consistent with the ‘early inhibition’ scenario (Fig. 1A) and suggesting that greater Elbow Rotation Decline may reflect premature anticipatory inhibition. Supporting this interpretation, no forearm lowering was observed in the imposed condition, where APA are unlikely.

The time interval 26 ± 15 ms before unloading was therefore identified as the optimal window for *Biceps brachii* inhibition during the voluntary unloading task. The ‘on-time’ *Biceps brachii* inhibition measure, hereafter referred to as EMG inhibition, was calculated as the average of preprocessed (see Methods) and baseline-corrected (-1 to -0.5 s) *Biceps brachii* EMG amplitude within a 26 ± 15 ms window before unloading. Importantly, the resulting EMG inhibition measure reflects two aspects simultaneously: the timing of *Biceps brachii* inhibition and its strength. Therefore, on individual trials, stronger EMG inhibition (more negative values) indicates both greater inhibition strength and optimal timing, whereas weaker EMG inhibition (more positive values) may reflect either reduced inhibition strength or its suboptimal timing, occurring too early or too late. However, since previous research suggests that the timing of *Biceps brachii* inhibition in adults is largely optimal, typically occurring within ∼50 ms before unloading with relatively low variability (Hugon et al., 1982; Viallet et al., 1987; Schmitz et al., 2002; Barlaam et al., 2012), we hypothesized that the EMG inhibition measure primarily reflects the strength of muscle inhibition.

EMG inhibition values, compared to elbow rotation parameters, were more normally distributed (Shapiro-Wilk test, p > 0.05 in 14 of 16 participants) and exhibited greater variability across trials within participants (mean ± SD: -0.018 ± 0.034; see Fig. 2B). A one-tailed one-sample t-test assessing whether the resulting EMG inhibition values were significantly negative across subjects indicated a significant decrease in EMG relative to zero (one-sided t-test: t(15) = -2.05, p = 0.029).

#### Reactive (long-latency) EMG inhibition during imposed unloading

In contrast to voluntary unloading, in the imposed condition, where load release triggered the unloading reflex, *Biceps brachii* suppression was time-locked to the unloading, resulting in a prominent reduction in EMG approximately 100 ms after load release in the group-averaged data (Fig. S2), spanning 82-117 ms (t_min_(15) = -6.20, p_min_ = 0.002). Reactive EMG inhibition, estimated as the mean EMG decrease over this time interval, was significantly below zero (one-sided t-test: t(15) = -4.35, p = 0.0003). This timing is consistent with previous reports of unloading reflex dynamics in the BLLT in adults (Dufossé et al., 1985; Schmitz et al., 2002; Barlaam et al., 2012) and falls within the range of long-latency EMG responses in upper-limb muscles (about 80-100 ms), which are thought to reflect task-dependent supraspinal modulations (Dietz et al., 1994).

### MEG data

#### EMG Inhibition correlates with high-gamma suppression in the medial SMA

To test whether anticipatory inhibition in the *Biceps brachii* is associated with decreased excitability in a cortical region controlling the muscle, we conducted a one-sample permutation cluster test against zero on the correlations between high-gamma power (averaged over 90-130 Hz), used as a proxy for regional excitability (Murthy and Fetz, 1996; Ray et al., 2008; Lundqvist et al., 2016; Riehle et al., 2018; Brazhnik et al., 2021), and EMG inhibition, across vertices of the right anticipatory motor network (see Supplementary Table S1). The analysis, conducted within the -0.25 to 0 s interval relative to unloading, revealed a significant correlation effect, peaking in the medial frontal cortex, specifically in the SMA (Fig. 3A), with the cluster spanning from -0.06 to -0.018 s (t_max_ = 6.41, p = 0.039, corrected). This finding suggests that stronger EMG inhibition 26 ± 15 ms before unloading is associated with a co-occurring reduction in high-gamma activity in the medial SMA.

In contrast to the voluntary unloading condition, no significant positive correlation was observed under the imposed unloading condition between high-gamma power and the strength of the unloading reflex (reactive EMG inhibition) in the *Biceps brachii* (t_max_(15) = 4.08, p_min_ = 0.68). See also the distribution of correlations between reactive EMG inhibition and high-gamma power (90-130 Hz) across both hemispheres in the Supplementary Results (Fig. S5) as well as the comparison of correlation coefficients between the voluntary and imposed unloading conditions, which showed a significant difference in the medial SMA (Fig. S6).

To further assess the potential contribution of other brain regions to anticipatory inhibition via alpha and beta activity, we computed Spearman’s rank correlations between periodic power (8-30 Hz) and EMG inhibition across sources in the right anticipatory motor network and subcortical structures (basal ganglia and cerebellum). The permutation cluster test revealed no significant correlation effect between EMG inhibition and alpha-beta band power in either cortical or subcortical regions.

#### SMA beta power correlates with EMG inhibition and high-gamma suppression

Transient increases in power caused by short-lasting oscillatory events – bursts – may obscure the relationship between periodic power and behavioral measures due to the complex non-linear dynamics of this activity (Lundqvist et al., 2024). We therefore hypothesized that, although no significant relationship between alpha or beta activity and EMG inhibition was observed at the right-hemisphere level, burst-like activity in these bands may still underlie the inhibition-related effects associated with the SMA – namely, stronger EMG inhibition and reduced high-gamma power, which might be observed at the label level. To test this possibility, we estimated the relationship between higher alpha- and beta-band periodic power and both stronger EMG inhibition and lower high-gamma power within the observed SMA cluster.

To examine whether anticipatory muscle inhibition was influenced by stronger alpha-or beta-band activity in the SMA, we computed Spearman’s rank correlations between SMA 8-30 Hz periodic power and EMG inhibition across time (-0.25 to 0 s relative to unloading). A one-sample, one-tailed permutation cluster test comparing correlations against zero revealed a significant negative correlation effect, peaking in the high-beta range (18-25 Hz) and spanning -0.228 to -0.018 s (Fig. 3B), suggesting that high-beta activity may support optimally timed anticipatory inhibition in the *Biceps brachii*.

EMG inhibition was therefore associated with both reduced high-gamma power and stronger high-beta power in the SMA. Notably, the temporal effect of late high-beta (24-25 Hz) power overlapped with the EMG inhibition interval and is consistent with the short latency between SMA stimulation and muscle response (mean MEP latency: 15.7 ms in Spieser et al., 2013; 22.6 ms in Entakli et al., 2014). We therefore hypothesized that stronger 24-25 Hz beta power may exert an inhibitory effect on SMA excitability, as reflected in reduced high-gamma activity, which aligns with prior evidence showing a negative relationship between beta and high-gamma/spiking activity (Ray et al., 2008; Lundqvist et al., 2016; Riehle et al., 2018). To test this, we computed partial Spearman’s correlation coefficients between SMA 24-25 Hz power and 90-130 Hz power, averaged over both frequency and the gamma correlation time window (-0.06 to -0.018 s; Fig. 3A), while controlling for broadband 1-130 Hz power to account for shared variance from broadband fluctuations (Spaak et al., 2012). This approach revealed significantly negative individual correlation coefficients (one-sample t-test: t(15) = -7.16, p < 0.001; Fig. S4B), indicating that stronger high-beta (24-25 Hz) power was associated with a more pronounced suppression of high-gamma (90-130 Hz) power, as predicted. A similar, albeit weaker, negative correlation was observed when periodic beta power (after removing the aperiodic component) was correlated with high-gamma power (one-sample t-test: t(15) = -2.17, p = 0.046). To test whether the observed beta-gamma correlation was specific to the high-beta range, we computed partial Spearman correlations between high-gamma power averaged over time and frequency (90-130 Hz; -0.06 to -0.018 s) and lower-frequency power across a broader time and frequency range (8-30 Hz; -0.25 to 0 s; Fig. S4A). As expected, significant correlations peaked at 25 Hz, but a secondary peak was observed at 9 Hz. However, because the 9 Hz effect was not linked to EMG inhibition (Fig. 3B), it likely reflects excitability fluctuations unrelated to anticipatory inhibition strength.

Since the relationship between EMG inhibition strength and reduced excitability was observed in the SMA, but not in M1, we further examined whether similar effects could be detected in the M1 elbow region. We examined correlations between EMG inhibition and both high-gamma power and periodic alpha-beta power (8-30 Hz). The M1 coordinates for the left elbow area (*Biceps & Triceps brachii*) were derived from Talairach space (Lotze et al., 2000; Plow et al., 2010), converted to MNI305 space, and averaged to yield [32, -20, 66] in the right hemisphere. The M1 elbow region was defined as the peak vertex and its six adjacent vertices, forming a seven-vertex M1 elbow label (see Fig. 3A, lower panel). The permutation cluster test did not reveal a significant correlation between EMG inhibition and either high-gamma power or 8-30 Hz periodic power in this region.

#### High-gamma suppression mediates the relationship between beta power and EMG inhibition

To explain the observed relationships between stronger EMG inhibition, reduced high-gamma power, and greater high-beta power in the SMA, we proposed a model in which the effect of high-beta activity on muscle inhibition is mediated by suppression of high-gamma activity. In this framework, beta activity is hypothesized to reduce SMA excitability, indexed by decreased high-gamma power, which in turn is linked to stronger anticipatory inhibition of the *Biceps brachii*. This model is consistent with prior evidence supporting an inhibitory role of beta activity on local excitability (Ray et al., 2008; Lundqvist et al., 2016; Riehle et al., 2018), and highlights a potential role of the SMA in controlling this muscle (Dum and Strick, 1991; Spieser et al., 2013; Entakli et al., 2014).

To test this model, we performed a mediation analysis examining whether the relationship between stronger high-beta (24-25 Hz) power and EMG inhibition was mediated by high-gamma (90-130 Hz) power reduction. Data were extracted from the SMA cluster source showing the strongest correlation between high-gamma power and EMG inhibition. In this model, beta power was the independent variable (X), EMG inhibition was the dependent variable (Y), and gamma power was the mediator (M). The GAMs, referred to as models A, B, C, and C’ (see Methods, *Mediation analysis* section), showed the following relationships: beta power predicted gamma power (*model A*: estimate (β) = -0.138, standard error (SE) = 0.027, 95% confidence interval (CI) = [-0.190,-0.085], t = -5.18, p < 0.001), gamma power predicted EMG inhibition (*model B*: β = 0.106, SE = 0.035, 95% CI = [0.037,0.175], t = 3.061, p = 0.002), beta power predicted EMG inhibition (*model C*: β = -0.069, SE = 0.025, 95% CI = [-0.120,-0.019], t = -2.725, p = 0.007), and beta power and gamma power together predicted EMG inhibition (*model C’*: beta power: β = -0.056, SE = 0.025, 95% CI = [-0.108,-0.005], t = -2.188, p = 0.029; gamma power: β = 0.092, SE = 0.035, 95% CI = [0.022,0.162], t = 2.609, p = 0.009). As expected, the four models showed significant associations among the tested variables. High-beta power was significantly associated with high-gamma power (model A), high-gamma power was significantly associated with EMG inhibition (model B), and high-beta power was also significantly associated with EMG inhibition (model C). When both predictors were included in the same model (model C’), high-beta and high-gamma power remained significantly associated with EMG inhibition, consistent with the proposed mediation framework.

The results of the mediation analysis are presented in Table 2 (see also Fig. 3C). The analysis revealed a significant mediation effect (Average Causal Mediation Effect, ACME), indicating that the association between high-beta (24-25 Hz) power and EMG inhibition was partly accounted for by its relationship with high-gamma (90-130 Hz) power. Specifically, this suggests that stronger high-beta power was associated with reduced high-gamma power, and that this association partly accounted for the observed relationship between stronger high-beta power and EMG inhibition. The total effect of high-beta power on EMG inhibition was also significant, reflecting the presence of an overall association between stronger high-beta power and more pronounced muscle inhibition, including both the direct relationship and the portion of the association mediated by high-gamma power. Importantly, the direct effect of high-beta power on EMG inhibition (Average Direct Effect, ADE) remained significant after accounting for the mediator, indicating that high-gamma suppression only partially explains the relationship between high-beta power and EMG inhibition. Consistent with this, approximately 18% of the high-beta–EMG inhibition association was mediated by high-gamma power. Together, these results support a partial mediation, whereby high-gamma suppression contributes to – but does not fully account for – the link between high-beta power and anticipatory muscle inhibition.

**Table 2.** Mediation analysis results showing that high-gamma power mediates the relationship between high-beta power and EMG inhibition. In this model, high-beta power (24-25 Hz) serves as the independent variable (X), EMG inhibition as the dependent variable (Y), and high-gamma power (90-130 Hz) as the mediator (M).

| | Estimate ($\beta$) | 95% CI | p-value |
| --- | --- | --- | --- |
| ACME | -0.013 | [-0.02, 0.00] | 0.006 ** |
| ADE | -0.057 | [-0.11, -0.01] | 0.027 * |
| Total effect | -0.069 | [-0.12, -0.02] | 0.007 ** |
| Proportion mediated | 0.183 | [0.04, 0.67] | 0.013 ** |

95% CI: 95% Confidence Interval; ACME: Average Causal Mediation Effect; ADE: Average Direct Effect.

### Beta burst analysis

Since beta activity in the brain typically manifests as brief bursts rather than sustained rhythms (Little et al., 2019; Lundqvist et al., 2024), we sought to validate our findings on beta-mediated inhibition with burst analysis. Specifically, we examined whether trials containing beta bursts exhibit anticipatory *Biceps brachii* inhibition and high-gamma power suppression.

Beta bursts (13-30 Hz) were detected in the SMA cluster vertex showing the strongest correlation between high-gamma power and EMG inhibition. Among the detected bursts, only those centered around the time and frequency range of the late correlation effect between high-beta power and EMG inhibition were selected for further analysis (-0.077 to - 0.018 s, and 22-28 Hz; Fig. 3B). This choice was motivated by the need to put constraints to select beta bursts potentially linked to anticipatory inhibition from those related to other processes. Considering the proposed mediation model (Fig. 3C), in which beta activity accounts for high-gamma suppression in the SMA, which in turn is linked to anticipatory *Biceps brachii* inhibition, we assumed that inhibition-related beta bursts should occur close to the gamma-to-EMG inhibition correlation effect (18-60 ms before unloading; Fig. 3A) given the high conduction velocity, and approximately 15-25 ms (according to Spieser et al., 2013; Entakli et al., 2014) before the muscle inhibition interval (26 ± 15 ms before unloading). Indeed, the 24-25 Hz beta power correlation with EMG inhibition (p < 0.05) spanning -77 to -18 ms before unloading (Fig. 3B) overlapped with the gamma correlation effect and reasonably preceded the muscle inhibition interval. We therefore constrained burst detection to this interval to capture only bursts that were closely time-locked to the inhibition-related effects. The frequency window was selected based on the peak frequency within the preselected time interval, which corresponded to 25 Hz. Considering the likely imprecision of standard time-frequency decomposition for asymmetric brain signals (Jones, 2016; Cole and Voytek, 2019), we allowed a broader frequency range for burst selection, spanning ±3 Hz around the 25 Hz peak. This selection also aimed to exclude the effect around 19 Hz (Fig. 3B), which lacked temporal precision, unlike the 24-25 Hz effect that was confined to the time of inhibition.

The percentage of trials containing the selected beta bursts was low, with a mean ± SD of 11.6 ± 1.9% per participant. The absolute number of trials with bursts was 10.1 ± 1.78 (mean ± SD). Notably, the number of trials with bursts remained consistent when the time-frequency signal was extracted from the 10 vertices of the medial SMA cluster and averaged for burst detection.

#### Beta bursts are linked to optimally timed EMG inhibition

The results above demonstrate that greater EMG inhibition is associated with increased high-beta power in the SMA. However, given that beta activity occurs in transient bursts, we suggested that the presence of bursts could more effectively predict the timing of anticipatory inhibition than beta power alone. To test this, we averaged baseline-corrected EMG amplitude across trials containing bursts for each subject. A permutation cluster test comparing this averaged EMG amplitude to zero revealed a significant decrease in EMG activity during burst trials, peaking at -0.026 s relative to unloading (t(15)=-3.92, p = 0.009; Fig. 4A), which coincides with the behaviourally defined optimal *Biceps brachii* inhibition window.

**Figure 4.**
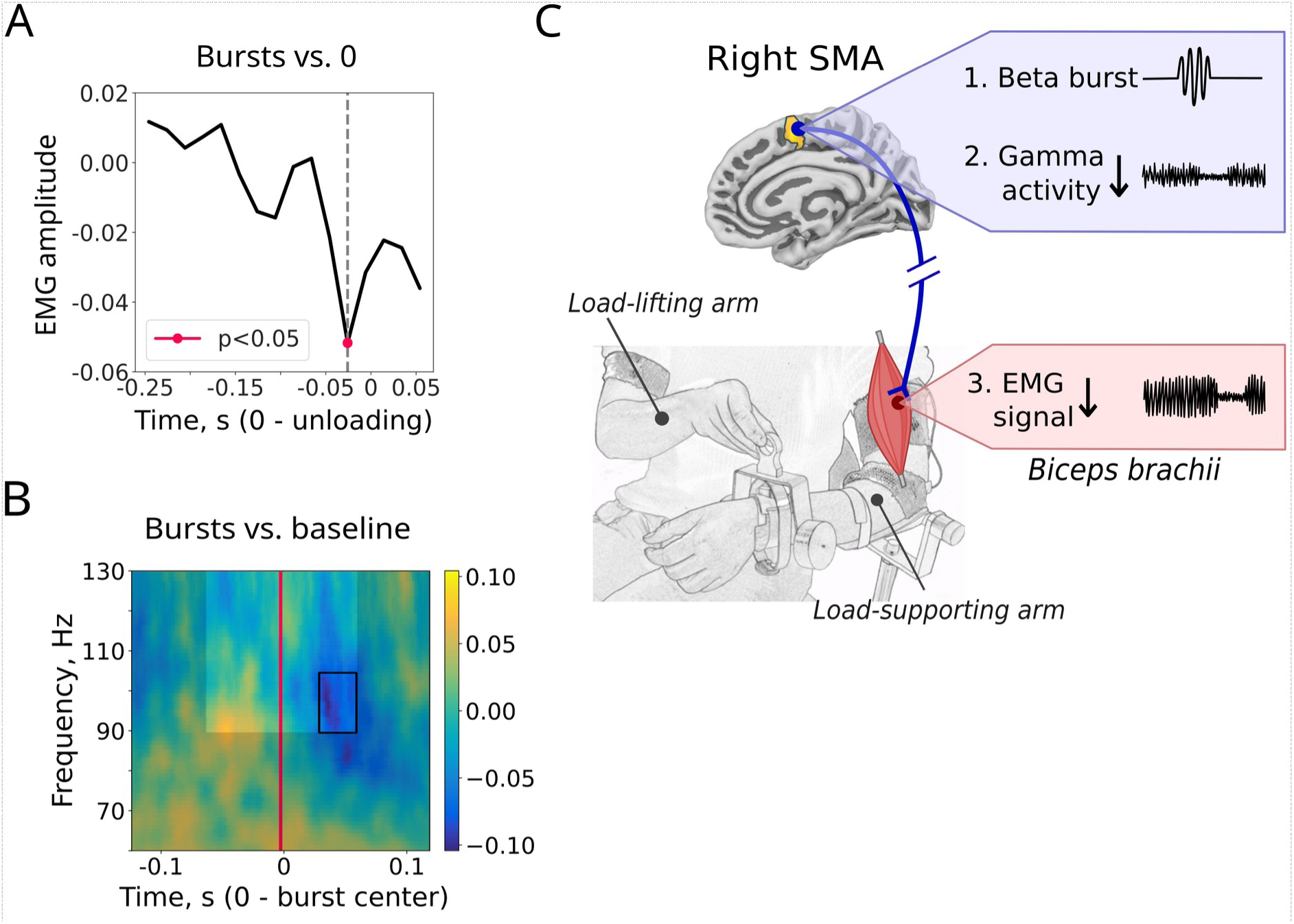
Beta bursts are linked to optimally timed *Biceps brachii* inhibition and high-gamma suppression. A hypothetical model is proposed linking SMA beta bursts, high-gamma suppression, and *Biceps brachii* inhibition. A: Baseline-corrected EMG amplitude averaged across SMA burst trials. Notably, the timing of the most pronounced EMG suppression coincides with the behaviourally defined optimal *Biceps brachii* inhibition window. **B:** Time-frequency contrast of gamma power between SMA burst and no-burst trials. Bright rectangle indicates the analysis window, and thin black lines mark time-frequency points showing significant contrast effects (p < 0.05, corrected). **C:** Schematic illustrating the hypothetical sequence of neural events during voluntary load-lifting leading to anticipatory inhibition of the postural elbow flexor, the *Biceps brachii*. An inhibitory beta burst (22-28 Hz) in the SMA contralateral to the muscle (1) leads to a reduction in excitability in this area, as reflected by suppression of high-gamma (90-130 Hz) activity (2), ultimately resulting in inhibition of the *Biceps brachii,* as reflected in EMG amplitude suppression (3). This cortical inhibition of the ongoing postural command associated with load support facilitates a smooth transition to a new postural state in anticipation of unloading, thereby stabilizing posture.

To test the specificity of the observed effect, we repeated the analysis using bursts selected within a broader 13-30 Hz range and within a lower-frequency 13-19 Hz range. A significant EMG decrease was observed for the 13-30 Hz range (t_min_ = -2.51, p_min_ = 0.036), whereas no significant effect was found for the 13-19 Hz range (t_min_ = -2.25, p_min_ = 0.089), suggesting that the effect observed for 13-30 Hz bursts is primarily driven by higher-frequency (22-28 Hz) bursts. We further tested whether the effect of 22-28 Hz bursts on EMG was specific to the selected time interval (-77 to -18 ms before unloading) by analyzing bursts within broader intervals of -100 to 0 ms and -150 to 0 ms. Bursts in the 22-28 Hz range selected within the -100 to 0 ms interval also showed a significant EMG decrease at the optimal time (t(15) = -3.28, p = 0.025). However, when a broader interval of -150 to 0 ms was used, the effect was reduced to a statistical trend (t(15) = -2.60, p = 0.088). These results suggest that the effect of higher-frequency (22-28 Hz) beta bursts on EMG aligns with the correlation effect (Fig. 3B), while not strictly limited to the selected intervals, supporting the sufficient robustness of the burst selection. Finally, to assess the robustness of beta burst effects to vertex selection, we normalized and averaged signals across the 10 vertices of the SMA label and extracted bursts within the same time-frequency range as in the main analysis (22-28 Hz; -77 to -18 ms relative to unloading). The EMG signal averaged across burst trials revealed a significant decrease specifically at the optimal time (t(15) = -2.92, p = 0.036), demonstrating that the observed effect is sufficiently reliable and not restricted to the peak SMA vertex.

Noteworthy, selecting an equal number of trials based on the highest beta power within the time-frequency window used for burst detection yielded no significant effect (p > 0.18), suggesting that the occurrence of a burst, rather than its amplitude, can better reflect its functional impact.

#### Beta bursts are linked to high-gamma power suppression

As described above, high-beta power negatively correlated with high-gamma activity in the SMA cluster region (see *SMA beta power correlates with EMG inhibition and high-gamma suppression* section and Fig. S4B). However, since beta activity occurs in bursts, we further tested whether the presence of beta bursts was associated with high-gamma power suppression within the same trials. A permutation cluster test confirmed a significant reduction in high-gamma power (90-130 Hz) in trials with beta bursts compared to baseline trials (Fig. 4B): in burst-centered data, high-gamma power decreased most prominently between 0.026 and 0.05 s following the burst center (t_max_ = -4.96, p_min_ = 0.032).

## Discussion

In this study, we investigated the neural mechanisms underlying anticipatory inhibition of the elbow flexor, the *Biceps brachii*, in the load-supporting arm, which counteracts forearm destabilization during voluntary load-lifting. Stronger, optimally timed muscle inhibition was associated with lower high-gamma power, potentially reflecting decreased cortical excitability, in the contralateral SMA, and to increased high-beta activity, with gamma suppression partially mediating this relationship. Notably, beta bursts (22-28 Hz) in the SMA were a stronger predictor of inhibition timing than beta power alone and were also associated with suppression of high-gamma activity. Together, our findings suggest that, in anticipation of load-lifting, high-frequency beta bursts may contribute to a reduction in high-gamma activity in the SMA, which in turn is associated with optimally timed inhibition of the contralateral *Biceps brachii* and stabilization of posture. To account for these observations, we proposed a physiologically plausible model (Fig. 4C).

### SMA, but not M1, mediates anticipatory Biceps brachii inhibition

During voluntary unloading, lifting a load induces a change in force, which is counteracted by timed anticipatory inhibition restoring equilibrium. The precise timing of this inhibition is essential for efficient bimanual coordination (Hugon et al., 1982; Viallet et al., 1987) and is altered in Parkinson’s disease (Viallet et al., 1987), Autism Spectrum Disorders (Schmitz et al., 2003), and other developmental motor-related conditions (Jover et al., 2006, 2010). In adults, anticipatory inhibition is closely time-locked to the same muscle activation in the load-lifting arm, suggesting a central timing mechanism coordinating both arms (Paulignan et al., 1989; Massion, 1992). However, developmental studies reveal that this precise temporal coordination is acquired gradually during childhood (Schmitz et al., 2002; Barlaam et al., 2012). Bolzoni et al. (2015) further demonstrated that APA can be selectively influenced through modulation of SMA activity without affecting voluntary movement. These findings suggest that APA are governed by a separate control mechanism, which integrates movement parameters and predicted postural disturbances to generate timely adjustments in muscle activity. To investigate the mechanism of anticipatory muscle inhibition, a core component of APA in the BLLT, we therefore focused on the right-hemispheric network contralateral to the postural arm where APA occur.

Although anticipatory inhibition is well-developed in adults, its latency still exhibits some variability (Barlaam et al., 2012). We observed that optimal forearm stabilization is achieved when *Biceps brachii* inhibition precedes the onset of unloading by 26 ± 15 ms, consistent with previously reported adult latencies (25 ms: Dufossé et al. (1985); 32 ms: Barlaam et al. (2012)).

Early studies have suggested that elbow flexor activity during the BLLT is controlled by the contralateral M1, and that M1 suppression may underlie anticipatory inhibition (Massion, 1992; Viallet et al., 1992; Kazennikov et al., 2005). This hypothesis was specifically tested by Kazennikov et al. (2005, 2006), who examined the relationship between elbow flexor activity and M1 excitability during APA. Using transcranial magnetic stimulation (TMS), they demonstrated reduced M1 excitability, reflected in decreased motor evoked potentials (MEPs), during anticipatory *Biceps brachii* inhibition. However, a similar MEP-EMG relationship was also observed under control conditions, suggesting that the effect was not specific to APA. In our study, we assessed high-gamma (90-130 Hz) power, which closely correlates with neural spiking (Ray et al., 2008; Lundqvist et al., 2016; Riehle et al., 2018; Brazhnik et al., 2021), as a proxy for excitability. Surprisingly, we also found no association between *Biceps brachii* inhibition and M1 excitability. However, stronger muscle inhibition was correlated with reduced excitability in the medial SMA – a relationship specific to voluntary unloading and APA, as it was not observed in the control condition, where load-release timing could not be anticipated.

The involvement of the SMA in BLLT has been consistently demonstrated in MEG and fMRI studies (Schmitz et al., 2005; Ng et al., 2011, 2013b). APA impairments following contralateral SMA lesions (Viallet et al., 1992) led authors to propose that the SMA gates postural stabilization circuits. In the present study, anticipatory *Biceps brachii* inhibition was linked to decreased high-gamma activity in the SMA, suggesting that this muscle inhibition may be associated with reduced SMA excitability. These findings support and extend previous accounts, suggesting that the SMA may contribute to the direct control of postural elbow flexors during voluntary load lifting. More specifically, our data are consistent with the view that, in anticipation of force changes, ongoing SMA activity controlling the postural *Biceps brachii* is suppressed – as reflected by reduced high-gamma activity – thereby enabling optimally timed muscle inhibition.

These results therefore support the previous speculations by Ng et al. (2011, 2013b) that the SMA, rather than M1, mediates APA via direct corticospinal projections. This interpretation is plausible, as the SMA displays precise somatotopic organization (He et al., 1995; Strother et al., 2012) and has corticospinal projections about half as extensive as those of M1 (Dum and Strick, 1991). Additionally, TMS stimulation of the SMA during motor tasks in humans evokes arm muscle MEPs similar in latency and amplitude to those from M1 (Spieser et al., 2013; Entakli et al., 2014). These findings indicate that the SMA can exert direct and precise control over arm muscles.

The assessment of neural excitability through high-gamma (90-130 Hz) activity during a motor task using a non-invasive technique may raise concerns about potential contamination from myogenic artifacts (Whitham et al., 2008; Muthukumaraswamy, 2013). We, however, argue that this is unlikely to affect our results. First, *Biceps brachii* inhibition was associated with decreased, rather than increased, high-gamma power, with the effect confined to the SMA on the medial surface, where tonic muscle contamination is expected to be low. Second, small head motion (<1 cm) and the absence of visible movement during the anticipatory period suggest minimal phasic EMG contamination. Combined with source localization based on spatial filtering, which reduces broadband myogenic gamma artifacts (Hipp and Siegel, 2013; Manyukhina et al., 2022), we argue that the observed correlation between muscle inhibition and high-gamma power reflects a genuine neural process.

In humans, SMA-originating corticospinal projections are more prominent than in non-human primates (Maier et al., 2002), and may have evolved to facilitate anticipatory control during complex hand movements (Chen et al., 2013). Such SMA-mediated hand muscle control, supported by beta-band coherence, has been demonstrated in tasks requiring precise, time-predictable force production in humans (Chen et al., 2013; Spieser et al., 2013; Entakli et al., 2014). In line with these findings, our results suggest that during voluntary load lifting, the SMA may exert partial, selective control over postural elbow flexors to ensure appropriately scaled APA. The release of this control – reflected by high-gamma suppression in the SMA – may be supported by beta bursts, facilitating a smooth transition to a new postural state. Whether this mechanism is specific to the optimal inhibition time interval, rather than earlier or later times, requires further investigation.

### Proposed mechanisms linking beta activity to anticipatory Biceps brachii inhibition

Beta-band activity has been extensively implicated in motor control, particularly through its established role in maintaining and adjusting muscle force production according to task demands (Peng et al., 2024; Lattari et al., 2010; Jana et al., 2020). Yet, the mechanisms by which beta activity mediates cortex-muscle interactions, ultimately shaping behavior, remain poorly understood (Khanna & Carmena, 2017; Peng et al., 2024). Although traditionally considered a sustained rhythm, increasing evidence indicates that beta activity is organized in transient oscillatory events – bursts (Lundqvist et al., 2024), and considering the true temporal structure of brain dynamics may advance our understanding of the neural mechanisms linking brain and muscle activity. In particular, cortical beta bursts have been shown to elicit time-locked, burst-like activity of similar duration and rate in contracting muscles (Bräcklein et al., 2022). Beta bursts have also been reported to establish synchrony between muscles during static postures (Simpson et al., 2024) and to promote cortico-muscular synchrony during sustained isometric muscle contractions (Echeverria-Altuna et al., 2022).

Another line of evidence links beta activity, and particularly beta bursts, to action slowing or cancellation: later burst timing is consistently associated with delayed responses (Khanna & Carmena, 2017; Little et al., 2019) and slower stopping (Hannah et al., 2020; Jana et al., 2020). These findings suggests that beta bursts may exert an inhibitory influence on muscles, yet the neural mechanisms underlying this effect remain unexplored. Our results align with this evidence and suggest a potential mechanism by which beta bursts contribute to inhibitory control over postural muscles during natural posture-movement coordination. Based on the observed relationships between anticipatory *Biceps brachii* inhibition, high-gamma activity, and beta bursts, we proposed a hypothetical model (Fig. 4C) in which, during load support, inhibitory beta bursts arising in the SMA shortly before load-lifting suppress ongoing SMA activity involved in postural muscle control. This suppression at the optimal time may release this control and thereby enable anticipatory postural muscle inhibition. This interpretation is consistent with the latency of the observed beta-to-muscle inhibition correlation in the SMA (∼20 ms; Fig. 3B), which closely matches previously reported MEP latencies following SMA stimulation (15.7 ms, Spieser et al. (2013); 22.6 ms, Entakli et al. (2014)). It is also supported by the mediation analysis, which showed that the relationship between beta activity and muscle inhibition was partly mediated by reduced high-gamma activity. Together, these findings suggest that beta-band activity contributes to anticipatory muscle inhibition by reducing SMA excitability, consistent with prior reports of an inverse relationship between beta and high-gamma/spiking activity in sensorimotor areas (Ray et al., 2008; Lundqvist et al., 2016; Riehle et al., 2018), corroborating their opposing functional roles – excitatory for high-gamma and inhibitory for beta (Ray and Maunsell, 2011; Lundqvist et al., 2024).

Alpha-band activity was another candidate for mediating anticipatory inhibition in the BLLT, however, we did not find evidence in favor of this. Previous studies linking alpha activity to motor control primarily describe its role in action withholding (Bönstrup et al., 2015; Köster and Meyer, 2023), consistent with the proposed “gating by inhibition” hypothesis for alpha oscillations (Jensen and Mazaheri, 2010). Alpha activity may therefore gate the selection of appropriate motor commands, rather than shaping the precision of ongoing muscle control, which is instead attributed to beta-band activity. Alternatively, while transient beta activity may mediate temporally precise, short-lived muscle inhibition, alpha activity, by creating a window of excitability, could influence the temporal profile of this inhibition lengthening its duration, or mediate longer-lasting muscle suppression, as proposed by Bonnefond and Jensen (2025).

### Network mechanisms of anticipatory inhibition

While our findings indicate that the SMA mediates the core events underlying anticipatory *Biceps brachii* inhibition during the BLLT, the circuitry conveying inhibitory bursts to the SMA remains unclear. We found no correlation between alpha-beta power and anticipatory inhibition in the examined cortical and subcortical right-hemispheric network. This suggests that inhibition strength may rely primarily on local SMA computations, whereas other regions may contribute to generating the APA command without directly influencing inhibition strength. Alternatively, burst-related dynamics may obscure such relationships, as transient fluctuations in power could prevent a reliable estimation of the link between beta signals and behavior (Lundqvist et al., 2024). Under these conditions, the occurrence of bursts may be a more accurate predictor of behavior than beta power itself, as demonstrated in our data (Fig. 4A) and in previous studies (Little et al., 2019; Enz et al., 2021). To address limitations related to the event-like nature of beta activity (Lundqvist et al., 2024), we analyzed high-gamma power (Fig. 4A) and performed a directed connectivity analysis (see Supplementary Materials for details; Figs. S7, S8) centered on SMA beta bursts. While limitations in spatial resolution and trial count prevent the connectivity analysis from conclusively identifying the full network underlying SMA beta bursts, this exploratory analysis suggests potential sources. Importantly, the results of this connectivity analysis were replicated in a subsequent study in children performing the BLLT (Manyukhina et al., 2026), suggesting that burst-centered connectivity analysis may provide reliable estimates for assessing transient synchrony. Future research would benefit from further investigation of the network underlying anticipatory inhibition, taking intrinsic neural dynamics into account to better elucidate the mechanisms governing its generation and timing.

### Limitations

One limitation of this study is the inability to track individual inhibitory events in the *Biceps brachii*, which may have limited the number of anticipatory inhibitions contributing to our analyses. In addition, the use of a predefined time interval for EMG inhibition assessment prevents dissociating whether the observed effects are primarily related to the optimal timing of anticipatory muscle inhibition, its strength, or both – a question that should be addressed in future research. Interpretation of marginally significant effects requires caution given the limited sample size of sixteen participants. This limitation further restricts our ability to draw firm conclusions regarding the involvement of subcortical structures – specifically the basal ganglia and cerebellum – in anticipatory inhibition, also owing to the lower signal-to-noise ratio of deep MEG sources (Attal & Schwartz, 2013). Moreover, the spatial resolution of MEG may be insufficient to accurately localize activity in small or closely adjacent cortical regions. Finally, our burst detection approach, based on predefined time-frequency windows, may not optimally capture the relationship between transient activity and behavioral measures. Since beta bursts are highly dynamic, such approaches may lead to attenuated or even misleading estimates (Lundqvist et al., 2024). Future work should therefore develop methods specifically tailored to transient oscillatory events, particularly to assess the link between bursts and behavioral measures, by taking into account the functional heterogeneity of beta bursts reflected in the variability of their waveforms and temporal dynamics (Szul et al., 2023). Such approaches would be particularly valuable for detecting transient network synchrony associated with distinct functional processes (Lundqvist et al., 2024).

## Conclusion

In conclusion, our findings support a cortical mechanism underlying anticipatory inhibition of the postural elbow flexors during the Bimanual Load-Lifting Task. Shortly before load lifting (∼80 ms), high-frequency (22-28 Hz) beta bursts in the SMA contralateral to the postural arm were linked to reduced high-gamma (90-130 Hz) activity, potentially reflecting decreased SMA excitability. The reduction in high-gamma activity was in turn associated with optimally timed inhibition of the postural elbow flexor, the *Biceps brachii*. Mediation analysis further showed that the relationship between high-beta (24-25 Hz) power and *Biceps brachii* inhibition is partially mediated by high-gamma suppression, supporting the proposed mechanism (Fig. 4C), in which beta bursts suppress ongoing SMA activity involved in postural muscle control, thereby releasing this control and enabling muscle inhibition. Together, these findings highlight a central role of the SMA in anticipatory postural control and suggest that beta bursts can contribute to inhibitory control at the level of muscle output by modulating cortical excitability.

## Acknowledgments

The authors would like to thank Jérôme Prado, Franck Lamberton, and Sarah Le Diagon for their valuable contributions to this study, as well as the staff of CERMEP–Imagerie du Vivant for their helpful assistance with data collection.

## Author contributions

V. Manyukhina: Data curation, Formal analysis, Methodology, Software, Validation, Visualization, Writing – original draft. O. Abdoun: Methodology, Software, Validation. F. Di Rienzo: Data curation, Investigation. F. Barlaam: Data curation, Investigation. S. Daligault: Data curation, Investigation. C. Delpuech: Methodology, Resources. M. Szul: Methodology. J. Bonaiuto: Methodology. M. Bonnefond: Conceptualization, Methodology, Project administration, Supervision. C. Schmitz: Conceptualization, Funding acquisition, Methodology, Project administration, Supervision. All authors contributed to Writing – review & editing.

