## Supplementary Materials for "Beta bursts in SMA mediate anticipatory muscle inhibition"

Abbreviated title: SMA beta bursts mediate anticipatory inhibition

Viktoriya Manyukhina[1]*, Oussama Abdoun[1], Franck Di Rienzo[1,2], Fanny Barlaam[1], Sébastien Daligault[3], Claude Delpuech[1,3], Maciej Szul[4,5,6], James Bonaiuto[4,5], Mathilde Bonnefond[1]†, Christina Schmitz[1]†

*corresponding author

†both authors equally contributed to the work

[1]Université Claude Bernard Lyon 1, CNRS, INSERM, Centre de Recherche en Neurosciences de Lyon CRNL U1028 UMR5292, COPHY, F-69500, Bron, France

[2]Université de Lyon, Université Claude Bernard Lyon 1, Laboratoire Interuniversitaire de Biologie de la Motricité-UR 7424, F-69100, Villeurbanne, France

[3]CERMEP-Imagerie du Vivant, MEG Departement, F-69500, Bron, France

[4]Université de Lyon, Université Claude Bernard Lyon 1, F-69100, Villeurbanne, France

[5]Institut des Sciences Cognitives Marc Jeannerod, CNRS, UMR5229, F-69500, Bron, France

[6]Department of Psychiatry and Psychotherapy, Faculty of Medicine, University of Tübingen, 72076 Tübingen, Germany

Corresponding author:

Viktoriya Manyukhina

Université Claude Bernard Lyon 1

43 Bd du 11 Novembre 1918, 69100, Villeurbanne, France

### Supplementary Methods

#### *Cortical segmentation and label selection*

| | Cortical parcellation labels from Glasser et al. (2016) |
|---|---|
| M1 | R_4_ROI |
| SMA | R_SCEF_ROI, R_6ma_ROI, R_6mp_ROI |
| PM | R_55b_ROI, R_6d_ROI, R_6a_ROI |
| CMA | R_24dd_ROI, R_24dv_ROI |
| PrCu | R_PCV_ROI, R_7Am_ROI, R_7Pm_ROI |
| SMar | R_PF_ROI, R_Pft_ROI, R_Pfop_ROI, R_Pfm_ROI, R_PFcm_ROI |
| dlPFC | R_8C_ROI, R_8Av_ROI, R_i6-8_ROI, R_s6-8_ROI, R_SFL_ROI, R_8BL_ROI, R_9p_ROI, R_9a_ROI, R_8Ad_ROI, R_p9-46v_ROI, R_a9-46v_ROI, R_46_ROI, R_9-46d_ROI |
| IFC* | R_44_ROI, R_45_ROI, R_IFJp_ROI, R_IFJa_ROI, R_IFSp_ROI, R_IFSa_ROI, R_47l_ROI, R_p47r_ROI |

**Table S1.** Regions of interest (ROIs) selected for MEG source-level analysis, representing key nodes of the right anticipatory motor network.

Primary motor cortex, M1; supplementary motor area, SMA; premotor cortex, PM; cingulate motor area, CMA; precuneus, PrCu; supramarginal gyrus, SMar; dorsolateral prefrontal cortex, dlPFC; inferior frontal cortex, IFC.

*Although IFC involvement was not observed in the BLLT task, we included this region in the connectivity analysis (see below) due to its well-established role in inhibitory motor control (Filevich et al., 2012; Xu et al., 2016).

##### *Estimation of Peak Elbow Rotation and Elbow Rotation Decline*

To estimate Peak Elbow Rotation and Elbow Rotation Decline in the voluntary unloading condition, elbow rotation time series were smoothed with a 5-point moving average (8.3 ms) and 1-point step (1.7 ms) to reduce noise. The time of maximal elbow deflection was then visually annotated in MNE-Python. Time series were subsequently epoched from -1.7 to 1.2 s relative to this point.

The amplitude of Elbow Rotation Decline was defined as the difference between (1) the average of the minimum values over a 33-ms window (excluding outliers) within the interval from 165 ms before unloading to the time of the annotated maximum elbow rotation, and (2) the average of the same number of maximal values (baseline interval) within the period from 165 ms before unloading to the last time point used for averaging the minimum values (this ensured exclusion of the subsequent positive elbow deflection). Therefore, in case of negligible elbow decline, time intervals for Elbow Rotation Decline and its baseline could overlap.

The amplitude of Peak Elbow Rotation was calculated as the difference between (1) the average of the maximal values over a 33-ms interval (excluding outliers) within a 165 ms window surrounding the annotated maximal elbow rotation, and (2) the average of the same number of minimum values (baseline interval) within the interval from 350 to 50 ms before unloading.

Elbow Rotation Decline and Peak Elbow Rotation were calculated individually for each trial. The resulting plots, including the time points used to estimate peak elbow rotation, elbow rotation decline, and their respective baselines, were visually inspected. Figure S1 presents the three example trials, the same as in the main manuscript (Fig. 1B), along with their corresponding analysis windows.

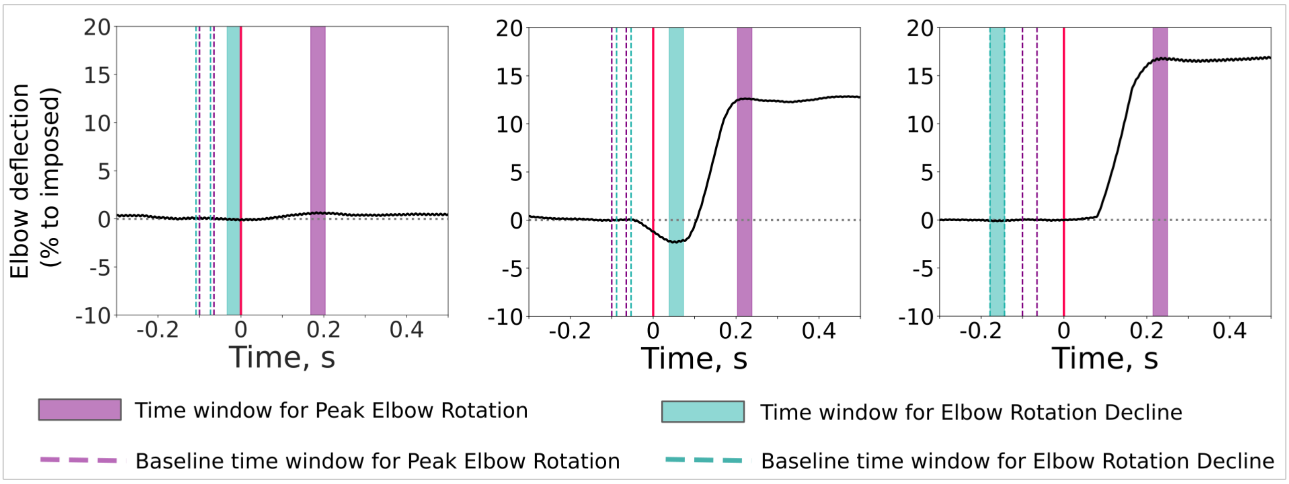


**Figure S1.** Single-trial elbow rotation amplitude examples aligned to unloading onset (time = 0), showing the time intervals used to estimate Peak Elbow Rotation and Elbow Rotation Decline. Values are presented as percentages of elbow deflection during voluntary unloading relative to the individual maximum, estimated as the median peak elbow rotation in the imposed condition.

#### *Evaluation of linearity assumptions in the mediation models*

As described in the main manuscript, a mediation analysis was conducted to determine whether the relationship between periodic beta power and EMG inhibition is mediated by high-gamma power, which is linked to both beta power and EMG inhibition. To do this, we verified linearity assumptions for the four linear models used (Table 1). First, the linearity of the relationships between the dependent and independent variables was visually inspected. The Durbin-Watson d-test (dwtest() function, 'lmtest' package, v. 0.9.40) was used to test for autocorrelation in the residuals of the estimated GAM models. The test indicated a significant positive autocorrelation for model a (DW = 1.89, p = 0.01). However, the DW value being close to 2 suggests that the autocorrelation is mild and may not substantially affect the validity of the model. For the other models, no autocorrelation was observed (mean DW ± SD: 1.970 ± 0.002, mean p-value ± SD: 0.163 ± 0.011). To assess the presence of multicollinearity among the linear predictors in models a and c', the

Variance Inflation Factor (VIF) was calculated (mgcv.helper::vif.gam() function, ‘mgcv.helper’ package, v. 0.1.9). All VIF values were < 2, suggesting an absence of multicollinearity.

##### *Evaluation of linearity assumptions in estimating the optimal inhibition time*

The analysis above revealed that stronger EMG decline around -0.026 s reflected the optimal timing for anticipatory inhibition, as indicated by greater forearm stabilization. To validate the use of linear regression models at this time, we tested whether the key assumptions were met for the subject-based models.

The linearity of the relationship between independent and dependent variables was confirmed through visual inspection of scatter plots. In all the subject-based models, the residuals were normally distributed (Shapiro-Wilk tests, $p > 0.05$ in all participants), did not show positive or negative autocorrelation supporting their independence (Durbin-Watson test, group-averaged statistics: 1.96 ± 0.19 (SD), min: 1.68, max: 2.36), and did not show the presence of heteroscedasticity (Breusch-Pagan test, $p > 0.05$ for all participants). To ensure that there was no multicollinearity, the Variance Inflation Factor (VIF) was calculated. Both independent variables had acceptable VIF values below 5, indicating that the predictors were not highly correlated (group-averaged VIF: 1.05 ± 0.05 (SD), min: 1.00, max: 1.19).

##### *Connectivity estimation*

As a complementary analysis, we investigated the right-hemispheric network contributing to the anticipatory inhibition. For this, we computed directed connectivity from a selection of right-hemispheric cortical regions associated with voluntary unloading (Table S1) to the medial SMA, which was linked to anticipatory inhibition strength. Specifically, we aimed to examine whether inhibitory beta bursts in the SMA receive directed influence in the beta band from other brain regions. We estimated connectivity in trials containing SMA beta bursts, aligning data to burst centers, and in an equal number of control trials, also aligned to burst centers for temporal shuffling. For directed connectivity estimation, we computed the Phase Slope Index (PSI) using the phase_slope_index() function in MNE-Python and validated the results using Granger causality, estimated with the spectral_connectivity_epochs() function. The choice of PSI over Granger causality for the main directed connectivity analysis was primarily motivated by its greater robustness to parameter choices in our dataset, together with its higher demonstrated resistance to mixed noise and its tendency to miss non-existent interactions rather than produce false positives (Nolte et al., 2008). We also considered that PSI-based connectivity estimated across the selected frequency band may help account for non-sinusoidal bursts (Szul et al., 2023), which can produce frequency modulations within a single burst and complicate source identification.

PSI and Granger causality were calculated from the vertices of the selected right-hemispheric cortical regions to the peak vertex of the SMA cluster, which showed the strongest correlation with *Biceps brachii* inhibition. Spectral decomposition was performed using Morlet wavelets (4 cycles). PSI was computed in the 22-28 Hz range (used for beta burst selection) separately for burst and no-burst control trials, and a PSI contrast between burst and no-burst trials was estimated. Granger causality was calculated for burst trials only, using 30 lags and the data rank, and averaged across the 22-28 Hz frequency range. For contrast estimation, Granger scores from burst trials were compared with scores computed in the opposite direction and with time-reversed Granger scores (Fig. S6B). A one-tailed spatio-temporal permutation cluster test was conducted within the -0.05 to 0 s window relative to burst centers on the resulting PSI and Granger causality contrasts. This window was chosen to include the time preceding the burst peak, allowing identification of brain regions whose activity reasonably precedes and may influence the SMA during its beta bursts. Additional tests examined contributions from the cerebellum and basal ganglia.

We hypothesized that regions functionally connected to the SMA during SMA beta bursts would also exhibit increased beta bursting in the same frequency range (22-28 Hz), assuming predominantly linear signal transmission. To test this, we assessed whether regions identified by PSI and corroborated by Granger causality showed elevated beta burst probability during SMA burst trials. Beta burst probability in these regions was calculated separately for SMA burst and no-burst control trials at each vertex, time and frequency point, defined as the proportion of trials containing bursts relative to the total number of trials. The probabilities were then averaged across vertices within each cluster and across frequencies. A one-tailed permutation cluster test was conducted to assess whether beta burst probability in PSI-defined regions was higher during SMA burst trials compared with no-burst trials, within the -0.05 to 0.05 s window.

### Supplementary Results

##### *Group-level Biceps brachii EMG suppression relative to load release*

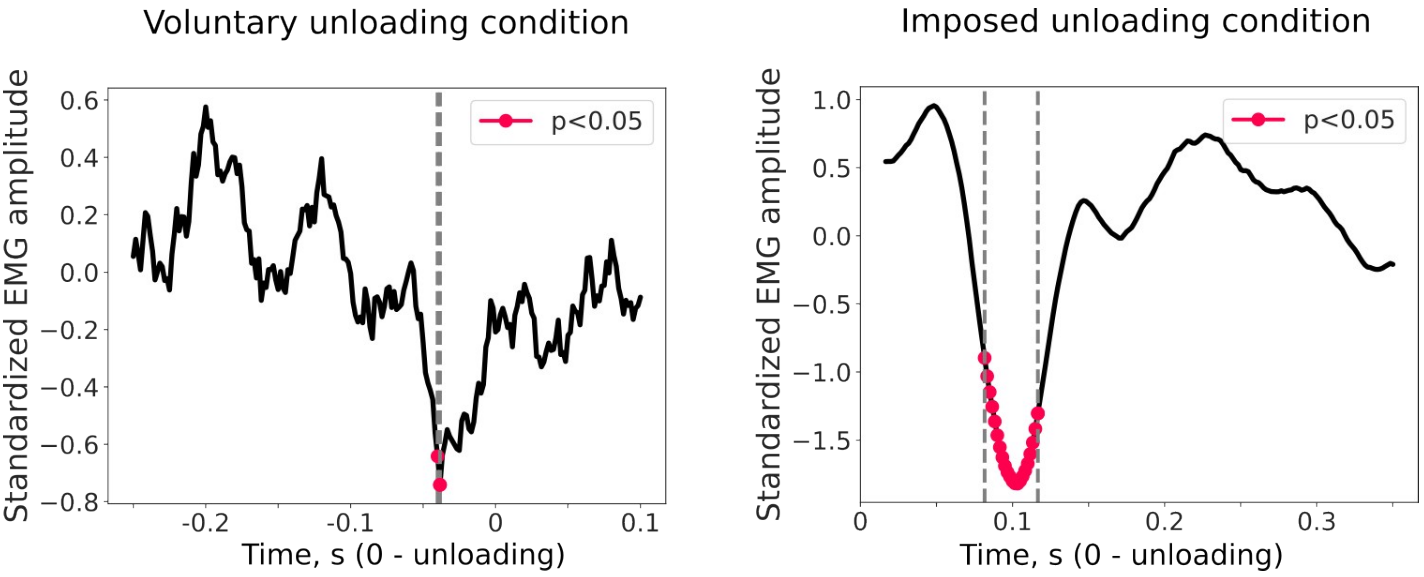


**Figure S2.** Group-averaged EMG amplitude showing time intervals with a significant decrease, corresponding to *Biceps brachii* inhibition, during voluntary unloading (left) and imposed unloading (right) ($p < 0.05$, TFCE-corrected).

##### *Interpretation of the link between EMG and elbow rotation across time*

To determine the optimal timing for anticipatory inhibition, we fitted linear regression models predicting *Biceps brachii* EMG from trial-by-trial variability in Peak Elbow Rotation and Elbow Rotation Decline over time (see Methods, *Estimation of the EMG inhibition* section). This analysis revealed three significant time intervals, -86± 15 ms, -66± 15 ms, and -26± 15 ms (Fig. S3, left panel), where EMG modulation was most strongly associated with elbow deflection.

However, closer examination of the correlation coefficients between EMG amplitude and the two elbow rotation metrics (Fig. S3, right panel) revealed that only around -26 ms was a stronger EMG suppression associated with both reduced (less negative) Elbow Rotation Decline and lower (less positive) Peak Elbow Rotation, indicative of a more stabilized forearm. This interval was therefore consistent with the ‘on-time’ inhibition scenario (Fig. 1A), and also corresponded to a reduction in group-averaged EMG activity relative to baseline (Fig. 2A, right panel).

By contrast, earlier time intervals -86± 15 ms and -66± 15 ms showed positive associations between EMG amplitude and Elbow Rotation Decline (Fig. S3, right panel), suggesting that stronger inhibition at these times was associated with forearm lowering, consistent with the ‘early inhibition’ scenario (Fig. 1A). Moreover, stronger Elbow Rotation Decline was significantly correlated with more pronounced *Biceps brachii* EMG suppression when measured specifically at -86± 15 ms and -66± 15 ms relative to unloading across trials (averaged over 30 ms; $t = 7.5$, $p < 0.001$ and $t = 5.2$, $p < 0.001$, respectively). These findings suggest that more pronounced Elbow Rotation Decline may reflect premature anticipatory inhibition within this paradigm. Supporting this interpretation, no forearm lowering was observed in the imposed condition, where APA were unlikely.

The present study, however, leaves open the question of whether these premature anticipatory *Biceps brachii* inhibitions are mediated by mechanisms similar to those underlying optimally timed inhibitions. Previous research suggests that adults predominantly exhibit optimally timed inhibitions (Schmitz et al., 2002; Barlaam et al., 2012). Consistent with this, no visible EMG decrease was observed at the group level within the premature inhibition interval (Fig. S2, left panel), suggesting that the number of such premature inhibitions was relatively small, limiting our ability to examine their underlying mechanisms in the current dataset. Future studies should examine these early anticipatory inhibitions using larger samples and sufficient trial numbers to reliably detect these events and assess their functional significance for anticipatory postural control.

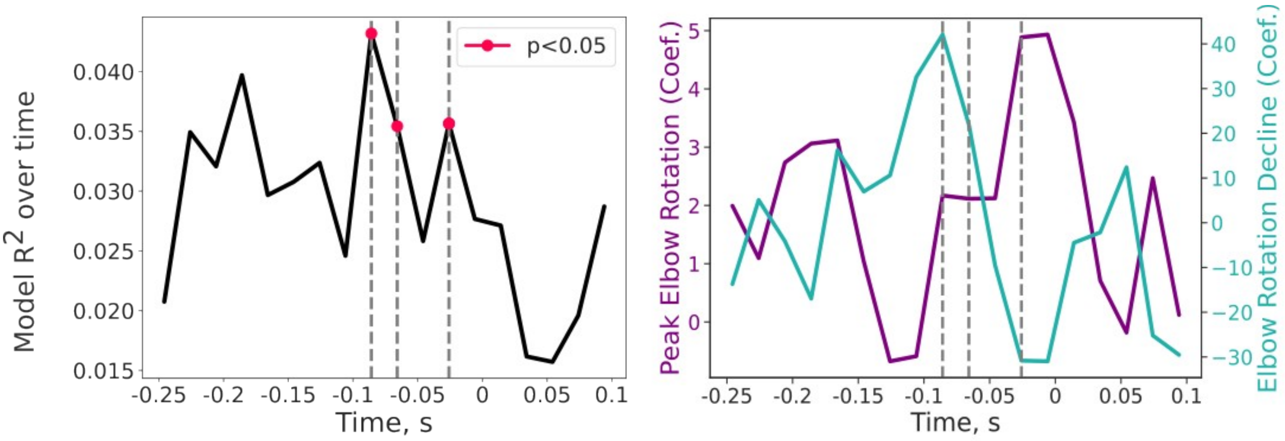


**Figure S3.** Parameters of linear regression models predicting EMG amplitude from elbow deflections. Data are aligned to unloading onset (time = 0). **Left panel:** Group-averaged $R^2$ values from linear regression models computed at each time point (dependent variable: *EMG amplitude*; independent variables: *Peak Elbow Rotation, Elbow Rotation Decline*); same as in the main manuscript, Figure 2C. **Right panel:** Group-averaged linear regression coefficients at each time point, provided separately for the two independent variables. Red markers and gray dashed lines denote time points where the relationship between EMG inhibition and the elbow deflection parameters reached significance.

##### *Correlation between 8-30 Hz power and high-gamma power*

To test whether the beta-to-gamma correlation was specific to both the gamma-to-EMG inhibition correlation interval and the high-beta range, suggesting that high-beta activity mediates SMA excitability linked to muscle inhibition, we computed a partial Spearman's correlation between high-gamma power (90-130 Hz; -0.06 to -0.018 s) averaged over frequency and time, and lower-frequency power across a broader time-frequency range (8-30 Hz; -0.25 to 0 s; Fig. S4A). A one-sample, one-tailed permutation cluster test comparing correlations against zero revealed a significant negative correlation effect, spanning all frequencies but peaking in the high-beta (25 Hz) and low-alpha (9 Hz) ranges. As expected, the timing of these peak effects overlapped with the gamma-to-EMG inhibition correlation interval (red dashed lines, Fig. S4A). While the effect around

25 Hz was predicted, the correlation between 9 Hz and high-gamma power was an unexpected finding. It is possible that low alpha power also modulates excitability alongside high-beta power, as demonstrated in previous studies in both visual and sensorimotor domains (Osipova et al., 2008; Haegens et al., 2011; Yanagisawa et al., 2012; Bonnefond and Jensen, 2015). However, since the 9 Hz effect was not associated with EMG inhibition (Fig. 4B), it may instead reflect excitability fluctuations unrelated to anticipatory inhibition strength.

It should be noted that this approach for correlating beta power with high-gamma power has limitations, as regressing out total spectral power may lead to overestimated negative correlation coefficients. Investigating the relationship between these two strongly positively correlated variables – due to their shared aperiodic baseline – is particularly challenging, especially since 90-130 Hz high-gamma power largely reflects aperiodic activity itself. Extracting the aperiodic component from beta power is not an optimal solution either, as it is difficult to ensure that no residual aperiodic activity remains within the periodic beta estimate. Nonetheless, we argue that the observed negative correlation between high-beta and high-gamma power reflects a genuine effect, since this relationship persists when using periodic beta power, in the generalized additive model (GAM) mediation analysis, and, critically, is consistent with the observed reduction in high-gamma power during high-frequency (22-28 Hz) beta bursts (see Results in the main manuscript).

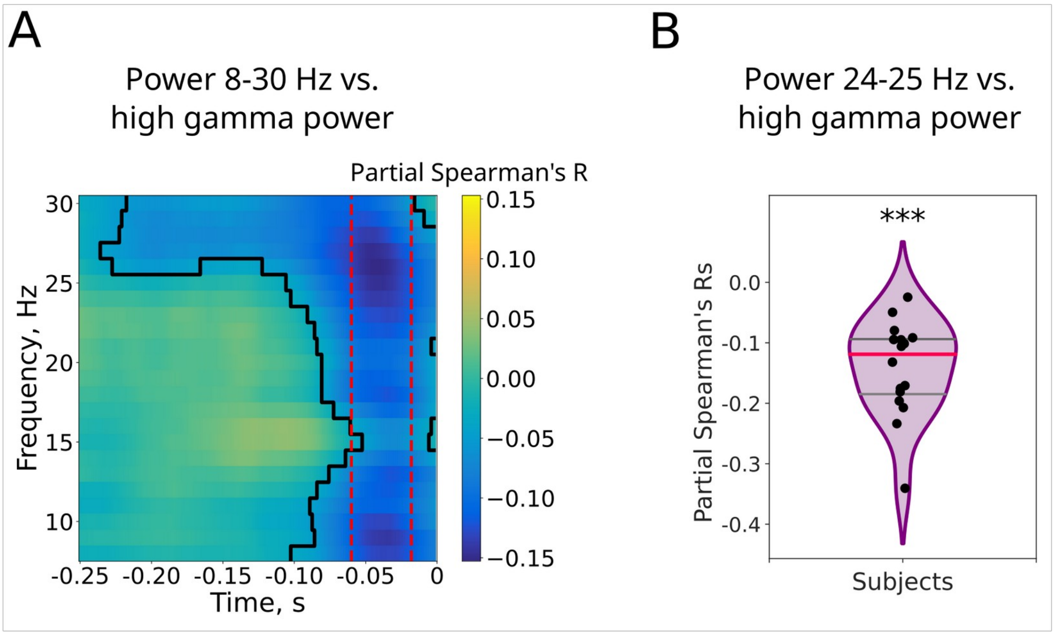


**Figure S4.** Relationship between 8-30 Hz power and high-gamma (90-130 Hz) power in the SMA cluster. **A:** Partial Spearman's rank correlation coefficients between total alpha-beta power and high-gamma power averaged over 90-130 Hz and the time interval corresponding to its correlation effect with EMG inhibition (indicated by red dashed lines). Data are aligned to unloading onset (time = 0). **B:** Violin plot showing individual subjects' partial Spearman's rank correlation coefficients between high-beta power (24-25 Hz) and high-gamma power (90-130 Hz), both averaged over the time interval of the gamma-to-EMG inhibition correlation effect.

##### *Relationship between reactive EMG inhibition and high-gamma power*

For demonstration purposes, we show the correlation between reactive EMG Inhibition and high-gamma power (90-130 Hz), used as an index of neural excitability, in the imposed condition across both hemispheres and from several viewing angles in Figure S5.

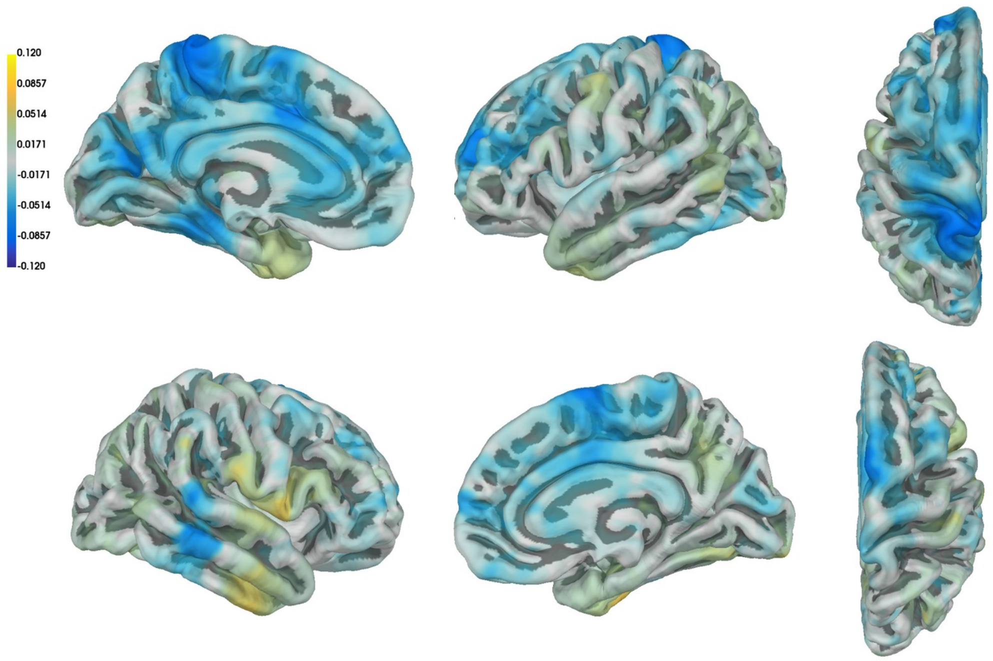


**Figure S5.** Correlation between reactive EMG inhibition and high-gamma power (90-130 Hz) across both hemispheres. No significant correlation was observed.

##### *Comparison of correlation coefficients between EMG inhibition and high-gamma activity in voluntary and imposed unloading*

To determine whether the observed correlation between EMG inhibition and high-gamma power in the voluntary unloading condition was specific to anticipatory inhibition, we compared these correlation coefficients with those obtained in the imposed unloading condition, where load release was unlikely to be anticipated. Because the temporal dynamics of *Biceps brachii* inhibition differed between voluntary and imposed unloading, we compared correlations averaged over time. In the voluntary unloading condition, we averaged the correlation coefficients over the time interval showing significant correlations (Fig. 3A). For the imposed condition, where no significant correlations were observed, we averaged the correlations within the interval from group-level *Biceps brachii* inhibition onset to its peak, assuming that high-gamma power modulation associated with inhibition should be strongest in this window (25-100 ms after unloading). The comparison of time-averaged correlation coefficients revealed a significant difference between conditions, with the maximal cluster (4 vertices) localized in the medial SMA and overlapping with the previously defined SMA cluster in which a correlation was observed in the voluntary condition ($t_{max}$ = 3.96, $p_{min}$ = 0.035; see Figure S6). The comparison between voluntary unloading and the control imposed condition yielded results comparable to those observed for voluntary unloading alone.

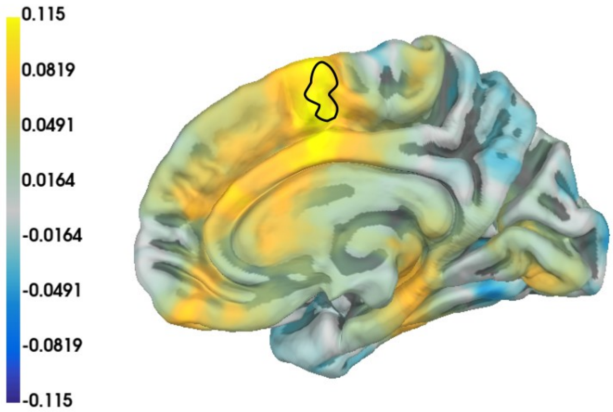


**Figure S6.** A cluster showing a significant difference in the correlation coefficients between EMG inhibition and high-gamma power when comparing voluntary unloading (averaged from -60 to -18 ms before unloading) and imposed unloading conditions (averaged from 25 to 100 ms after unloading).

##### *Connectivity analysis results*

To identify brain regions contributing to anticipatory inhibition, specifically those communicating with the SMA during inhibitory beta bursts and potentially involved in generating these signals, we conducted a directed connectivity analysis using the PSI and validated the results with Granger causality.

A one-tailed permutation cluster test revealed a significant PSI difference between burst and no-burst trials, with two key clusters identified: one near the M1 elbow representation (p = 0.043) and one in the lateral PFC (lPFC) (p = 0.004; Fig. S7A), most pronounced in the 50 ms preceding the SMA burst center (Fig. S7B). Granger causality analysis revealed similar, though not identical, clusters (Fig. S8A), including M1 (p = 0.044), lPFC (p = 0.057), and premotor cortex (PMC; p = 0.083). No significant differences were detected in the right basal ganglia or cerebellum in either analysis.

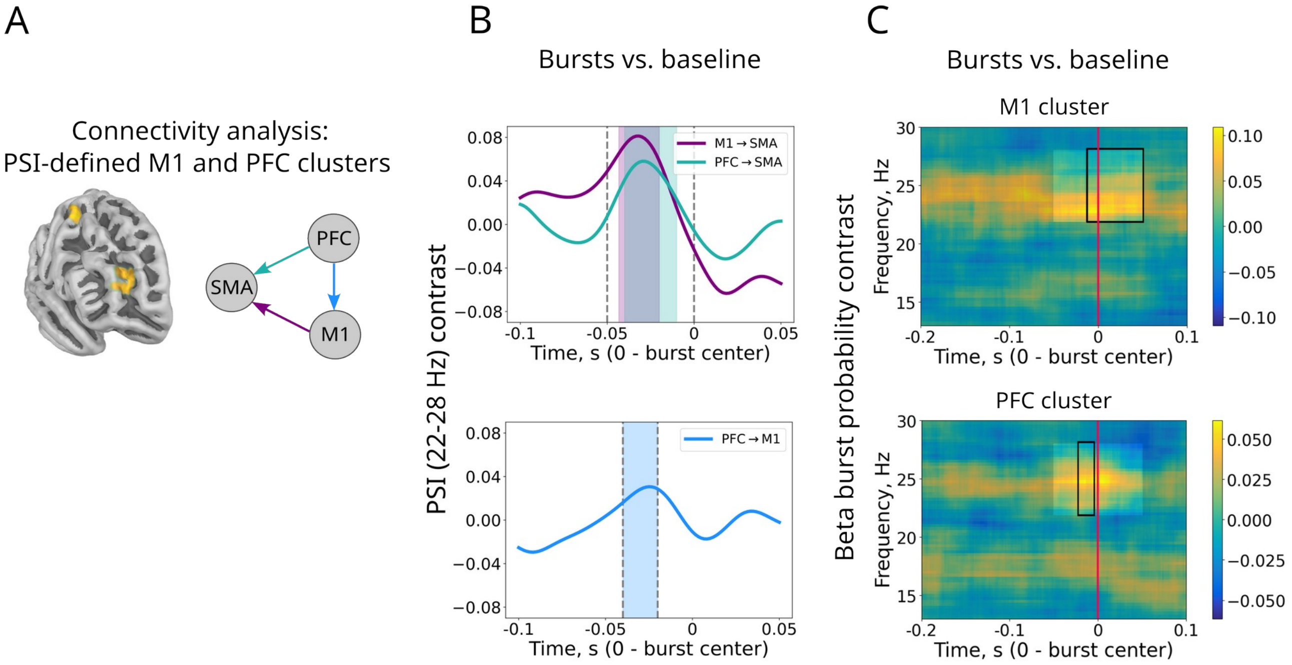


**Figure S7. A:** PSI-derived connectivity cluster regions (M1: 3 vertices, MNI305 peak: [38,-8,56]; lateral PFC: 10 vertices, MNI305 peak: [31,47,11]) exhibiting increased directed connectivity toward the SMA cluster during SMA beta burst trials relative to no-burst trials. Connectivity directions are indicated by schematic arrows. **B:** PSI timecourses for M1-to-SMA, PFC-to-SMA, and PFC-to-M1 contrasting SMA burst and no-burst trials. In the upper panel, dashed lines outline the analysis window, with shaded areas highlighting time intervals of the most prominent effects ($p < 0.05$, corrected). The lower panel marks the overlapping period of these effects, within which the average PSI contrast was significantly different from zero. **C:** Time-frequency contrast of beta burst probability between SMA beta burst and no-burst trials in the M1 and PFC cluster regions. Note that for this analysis, burst probability was averaged within the 22-28 Hz frequency range. Bright rectangles indicate the analysis window, and thin black lines mark time points showing significant contrast effects ($p < 0.05$, corrected).

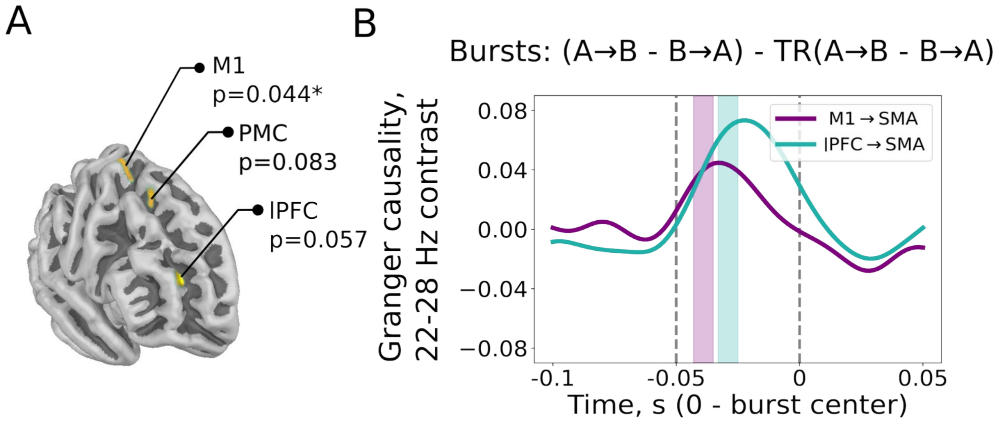


**Figure S8. A:** Granger causality-derived connectivity cluster regions exhibiting increased directed connectivity toward the SMA cluster during SMA burst trials. The Granger scores were contrasted with those computed in the opposite direction and with those obtained from time-reversed (TR) series. **B:** Granger causality timecourses for M1-to-SMA and PFC-to-SMA, contrasted with scores in the opposite direction and those from time-reversed signals. Dashed lines outline the analysis window, with shaded areas highlighting time intervals of the most prominent cluster effects ($p < 0.05$, corrected).

To investigate the interaction between the lPFC and M1 clusters during their concurrent influence on the SMA, we computed PSI between these regions and contrasted burst versus no-burst trials across vertices. The PSI contrast was then averaged across the vertices and the time interval corresponding to their overlapping effect on the SMA. Comparing the PSI contrast to zero across subjects revealed significant directed connectivity from the lPFC to the M1 cluster (one-sample t-test, $t = 2.33$, $p = 0.034$; Fig. S7B, lower panel).

To assess whether regions functionally connected to the SMA exhibit increased beta bursting during SMA bursts, we examined burst probability in the PFC and M1 clusters, averaged within the 22-28 Hz range in which connectivity was estimated. A permutation cluster test confirmed that both M1 and PFC clusters displayed higher burst probability in SMA burst trials compared to no-burst trials, with the effect most pronounced around the SMA burst centers (Fig. S7C).

The analyses above suggest that during inhibitory beta bursts in the SMA, the M1 elbow area and lPFC exert directed influence on the SMA in the burst frequency band, a finding further supported by increased burst probability in both regions during SMA bursts. However, as neither region is part of the action cancellation network (Borgomaneri et al., 2020), it is unlikely that either the lPFC or M1 generates the inhibitory bursts transmitted to the SMA; the source of these bursts therefore remains unclear. M1 may provide proprioceptive feedback to the SMA (Nasrallah et al., 2019) or relay a copy of the motor command from the left hemisphere (Viallet et al., 1992). The identified lPFC subregion, BA46, which exerts a facilitatory influence on M1 and modulates motor control in a task- and muscle-specific manner (Buschman et al., 2012; Hasan et al., 2013; Khan et al., 2024), may help maintain optimal beta synchrony in the SMA, thereby supporting anticipatory muscle inhibition.

One potential candidate for transmitting the inhibitory beta signal to the SMA is the pre-SMA. Unlike the SMA-proper, which is somatotopically organized, projects to the spinal cord, and connects with M1 (Tanji, 1994; Coull et al., 2016), the pre-SMA, located more anteriorly, maintains strong connections with the PFC, particularly BA46, and, together with the IFC, constitutes a core node of the action-stopping network, selectively suppressing motor activation according to task demands (Burle et al., 2004; Carbonnell et al., 2013). It is therefore plausible that the pre-SMA transmits timed inhibitory beta signals to the SMA-proper, suppressing the *Biceps brachii*. These beta bursts may propagate directly via adjacent arm fields (Luppino et al., 1993) or indirectly through the basal ganglia (Wadsley et al., 2022). Future research is needed to clarify the origin of inhibitory beta bursts mediating anticipatory *Biceps brachii* inhibition in the BLLT.

Estimating connectivity in brain signals characterized by bursts presents a particular challenge, as it must capture transient network synchrony rather than sustained communication, which standard measures are likely to miss (Lundqvist et al., 2024). Here, we partially addressed this issue by focusing our analysis on bursts of interest and restricting connectivity estimation to the surrounding time windows. However, the limited number of trials in which beta bursts within the predefined time-frequency range were detected (~10% of all trials) prevents us from drawing definitive conclusions and highlights the need for further investigation of the network supporting anticipatory inhibition. Two main limitations of this approach should be addressed in future research. First, the time-frequency window for burst selection was defined based on correlation results, which is suboptimal due to spurious spectral peaks from burst waveform variability (Jones, 2016). While our burst selection approach was supported by the EMG decline observed at the optimal time in trials with bursts (Fig. 4A), different bursts may reflect distinct processes, and those unrelated to anticipatory inhibition could, by chance, fall within the selected interval, or relevant bursts could be excluded, introducing noise into the estimation of transient network synchrony. Second, our connectivity analysis specifically focused on the high-beta frequency range, associated with *Biceps brachii* inhibition in the present study, and thus did not capture potential nonlinear network interactions likely induced by bursts (Lundqvist et al., 2024). Further research is therefore needed to clarify which brain regions, beyond the SMA, contribute to anticipatory muscle inhibition and how this network interacts with left-hemisphere activity related to load lifting.

##### *Control analysis of EMG oscillatory activity*

Apparent fluctuations in EMG amplitude preceding unloading (Fig. S2, left panel) may resemble oscillatory activity, suggesting that muscle inhibition follows EMG oscillations. However, this is unlikely. Previous studies using trial-averaged BLLT data did not report such fluctuations (Ng et al., 2011, 2013a, 2013b), and single-trial analyses consistently reported a single anticipatory inhibition (Hugon et al., 1982; Schmitz et al., 2002; Barlaam et al., 2012). Similarly, in our data, the group-averaged EMG decrease occurred only within the time window reported in the literature (Fig. S2), suggesting that the apparent oscillations reflect noise.

To assess whether these fluctuations reflect EMG oscillatory activity, we performed a time-frequency analysis of EMG trials using Morlet wavelets (3 cycles) in the -1 to 0.5 s interval relative to unloading. Power was log10-transformed and baseline-corrected using the -0.75 to -0.5 s interval. A one-sample permutation cluster test in the -0.25 to 0.1 s time window and 8-30 Hz frequency range revealed no significant increase in power ($t_{max}$ = 2.72, $p_{min}$ = 0.25; Fig. S9).

Around the time of the EMG decrease (Fig. S2, left panel), the effect direction was negative (Fig. S9), indicating reduced beta-band activity during anticipatory inhibition, whereas the pre-inhibition period showed a positive change. This suggests that the EMG inhibition at -26 ± 15 ms and the preceding fluctuations reflect distinct processes. Overall, there is no evidence for increased alpha-beta periodic activity, supporting the interpretation that the observed fluctuations reflect noise rather than true oscillations.

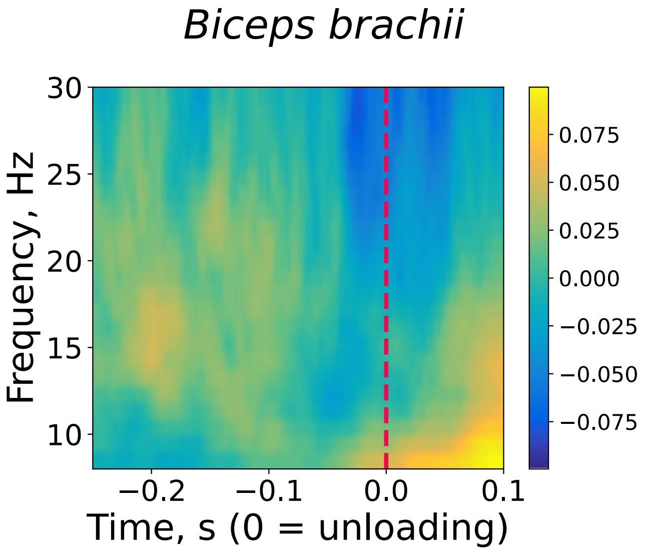


**Figure S9.** Group-averaged time-frequency power (baseline-corrected) estimated from the left *Biceps brachii*. No significant power increase was observed.

##### ***Reactive EMG inhibition in the imposed condition: Biceps brachii and Brachioradialis***

Voluntary unloading elicits anticipatory inhibition in the two elbow flexors that ensure load support: the *Biceps brachii* and *Brachioradialis*. In this study, we focused on the *Biceps brachii*, as the *Brachioradialis* exhibited a lower signal-to-noise ratio. Unlike the *Biceps brachii*, which showed a clear EMG suppression in this condition (~100 ms; Fig. S10, right), the *Brachioradialis* decrease was not significant, though a trend was observed ($t_{min}$ = -3.67, $p_{min}$ = 0.05; Fig. S10, left). We therefore concluded that *Brachioradialis* signal was unreliable for analysis in the voluntary unloading condition.

However, given that anticipatory inhibition in the *Biceps brachii* and *Brachioradialis* occurs within the same trials with comparable durations (92 ms vs. 82 ms) and onset times (-14 ms vs. -8 ms; Barlaam et al., 2011), their underlying mechanisms may be similar. Future work should reliably detect single-trial inhibitory events to determine whether inhibition of both flexors, and *Triceps brachii* activation, as reported in children (Schmitz et al., 2002), share the same SMA-mediated control.

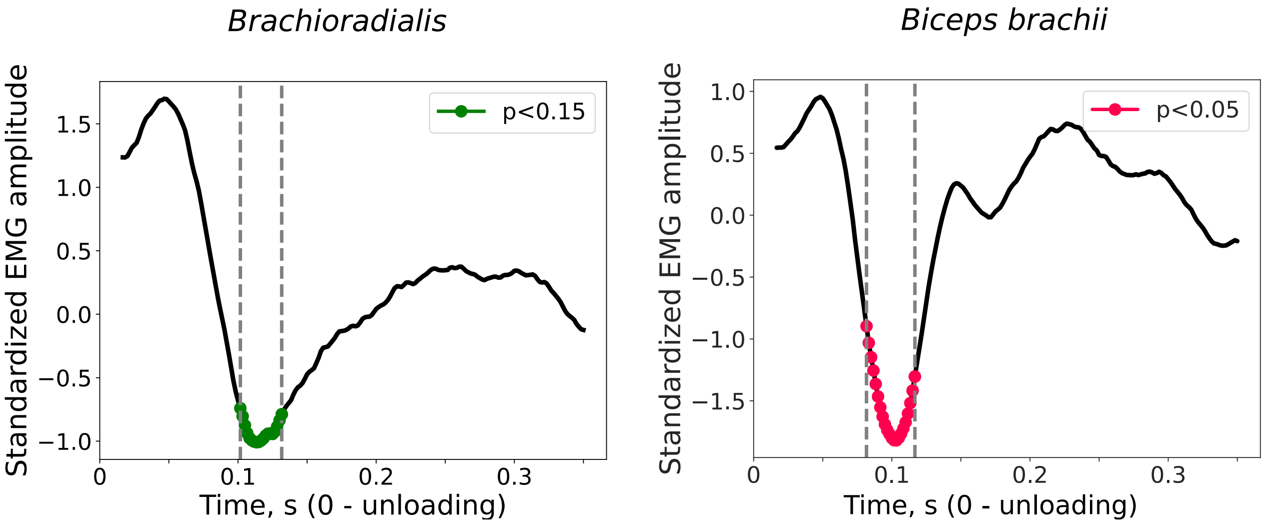


**Figure S10.** Group-averaged EMG amplitude modulation for the left *Brachioradialis* (left) and left *Biceps brachii* (right) during imposed unloading. Red points indicate the time interval with a significant decrease. Note that no significant decrease was observed in the *Brachioradialis* in contrast to the *Biceps brachii* ($p < 0.05$, TFCE-corrected)
